# Liquid-liquid phase separation and aggregation of the prion protein globular domain modulated by a high-affinity DNA aptamer

**DOI:** 10.1101/659037

**Authors:** Carolina O. Matos, Yulli M. Passos, Mariana J. do Amaral, Bruno Macedo, Matheus Tempone, Ohanna C. L. Bezerra, Milton O. Moraes, Marcius S. Almeida, Gerald Weber, Sotiris Missailidis, Jerson L. Silva, Anderson S. Pinheiro, Yraima Cordeiro

## Abstract

Structural conversion of cellular prion protein (PrP^C^) into scrapie PrP (PrP^Sc^) and subsequent aggregation are key events for the onset of Transmissible Spongiform Encephalopathies (TSEs). Experimental evidences support the role of nucleic acids (NAs) in assisting the protein conversion process. Here, we used the SELEX methodology to identify two 25-mer DNA aptamers against the globular domain of recombinant murine PrP (rPrP^90-231^), namely A1 and A2. High-affinity binding of A1 and A2 to rPrP was verified by ITC. Aptamers structure was characterized by theoretical predictions, CD, NMR and SAXS, revealing that A1 adopts a hairpin conformation. Aptamer binding caused dynamic aggregation of rPrP^90-231^, resulting from the ability of rPrP^90-231^ to undergo liquid-liquid phase separation (LLPS). While free rPrP^90-231^ phase separated into large droplets, aptamer binding increased the amount but reduced the size of the condensates. Strikingly, a modified A1 aptamer that does not adopt a hairpin structure induced transition to an ordered state, suggestive of amyloid formation on the surface of the droplets. Our results describe for the first time PrP:NA interaction leading to LLPS and modulation of this effect depending on NA structure and binding stoichiometry, shedding light on the role of NAs in PrP misfolding and TSEs.

## INTRODUCTION

Prion protein (PrP) is a ubiquitous glycosylphosphatidylinositol (GPI)-linked glycoprotein, highly conserved in mammalian species (1, 2). The conversion of the cellular prion protein (PrP^C^) into a pathological form, the scrapie PrP (PrP^Sc^), leads to prion diseases or Transmissible Spongiform Encephalopathies (TSEs). TSEs, such as Creutzfeldt-Jakob disease in humans (3, 4), scrapie in sheep (5), and bovine spongiform encephalopathy in cattle (6) are invariably fatal neurodegenerative disorders characterized by accumulation of PrP^Sc^ in the central nervous system and the precise mechanism of PrP^c^ conversion into PrP^Sc^ remains unknown (7).

Mature PrP^C^ is monomeric, susceptible to digestion by proteinase K, and harbors an intrinsically disordered N-terminal domain (residues 23 to ~121) as well as a globular C-terminal domain (residues ~121-231) enriched in α-helices. In contrast, PrP^Sc^ is mainly formed by β-sheets, is insoluble in aqueous solution, and aggregates into species resistant to proteolytic digestion (8–10). Although the protein-only hypothesis postulates that PrP^Sc^ is the solely factor responsible for TSE development (11), PrP^C^ to PrP^Sc^ *in vitro*-conversion assays allowed identification of additional cofactors that may participate in this process (12, 13). Among such cofactors, polyanionic molecules, including nucleic acids (NAs), were described to catalyze the conversion of PrP^C^ into PrP^Sc^-like species (13–15), possibly by lowering the energy barrier between these two PrP conformations (14). PrP can interact with both DNA and RNA molecules and such interactions were described to occur *in vitro* and *in vivo* (14, 16). Different binding affinities and stoichiometries were observed for PrP:NA complexes, depending on the NA size and sequence and on the PrP construct (17, 18). Moreover, the effect of NA binding on the structure, oligomerization profile and toxicity of the resultant PrP species to mammalian cells in culture are also variable (19–21). Previous reports suggest that DNA ligands with high GC content mediate loss of native secondary structure in recombinant PrP (rPrP), leading to protein aggregation into a cytotoxic species (21). In addition, a G-quadruplex DNA binds to rPrP with nanomolar affinity, leading to reciprocal conformational changes in both protein and NA (22), suggesting that the DNA secondary structure plays a central role in PrP binding. Three regions along full-length PrP (PrP^23-231^) were assigned to bind NAs: the first (residues 23-27) and second (residues 101-110) lysine clusters within the N-terminal domain have been shown to interact with both DNA and RNA, while the C-terminal globular domain (residues ~121-231) has been implicated in DNA binding (19, 22–24). Thus, based on the current literature, it is still debatable whether PrP:NA binding specificity and biological outcome are driven by the DNA/RNA sequence, structure and/or size. Moreover, binding stoichiometry seems to be crucial for PrP oligomerization induced by NAs (14, 25), as low protein to NA ratios increase aggregation.

Other studies have further explored the NA binding ability of PrP to select and characterize NA sequences capable of binding to PrP^C^ or the scrapie form with high affinity and specificity (17, 26–28), being valuable for therapeutic or diagnostic approaches. These sequences, named aptamers, were mostly identified by SELEX (<u>S</u>ystematic <u>E</u>volution of <u>L</u>igands by <u>Ex</u>ponential Enrichment) (17). Some of the identified NA aptamers significantly reduced PrP^Sc^ formation in cell-free assays; aptamers that tightly bind to and stabilize PrP^C^ are expected to block conversion and, thereby, prevent prion diseases (17, 26, 27, 29). Other aptameric sequences were described to bind β-sheet-enriched PrP structures, i.e. scrapie-like forms, with application in the diagnostic of TSEs (17).

Despite the body of work on the subject, there is still lack of high-resolution structural information about the PrP:NA complexes and there are many open questions about the biological effects of PrP:NA interaction. Here, we used SELEX to select two 25-mer single-stranded DNA aptamers (herein referred as A1 and A2) against the globular domain of murine recombinant PrP (rPrP^90-231^). Isothermal titration calorimetry (ITC) revealed higher binding affinity against rPrP^90-231^ for A1 (nanomolar) than A2 sequence (micromolar), which may be related to the ability of A1 to form a hairpin. Along our attempts to stabilize soluble rPrP:DNA complexes, we observed that both aptamers induced loss of protein secondary structure, either by aggregation and/or partial unfolding, in a stoichiometry-dependent manner. Furthermore, our scattering and microscopy data indicated dynamic formation of higher order assemblies, such as biomolecular condensates, immediately upon aptamer addition. Considering that reversible liquid-liquid phase separation (LLPS) is a hallmark of RNA-binding proteins, including proteins related to degenerative disorders (25, 30, 31), we investigated whether A1 and A2 could induce rPrP^90-231^ LLPS. In addition, it was recently described that full-length PrP can undergo phase separation induced by Amyloid-β oligomers (32). We observed that rPrP^90-231^ phase separates in aqueous solution into large droplets and that aptamer binding increases the amount but reduces the size of the protein-rich condensates. The aptamer effect on rPrP^90-231^ LLPS was clearly dependent on the protein:NA molar ratio, as observed for other proteins that drive phase separation (31). A modified A1 aptamer (A1_mut) that does not adopt a hairpin structure induced formation of a less dynamic, aggregation-prone material, suggestive of amyloid formation on the surface of liquid droplets. This liquid-to-solid state transition is a characteristic feature of other IDPs involved in neurodegenerative diseases, such as FUS and TDP-43 (33, 34). Our results shed light on the role of NAs in prion protein misfolding and describe for the first time PrP:NA interaction leading to LLPS and modulation of this effect depending on NA structure and binding stoichiometry.

## MATERIAL AND METHODS

### Expression and Purification of recombinant PrP

Both full-length (rPrP^23-231^) and the globular domain (rPrP^90-231^) of recombinant murine prion protein were expressed in *Escherichia coli* BL21 (DE3) cells and purified by nickel-affinity chromatography and in-column refolding, as described previously (35). The N-terminal six-histidine tag was not removed from rPrP^90-231^ for column immobilization in the SELEX procedure.

### SELEX

Aptamer DNA library was obtained from Euzoia Limited (United Kingdom). All oligonucleotides were flanked by primers: forward (5’–GGGAGACAAGAATAAACGCTCAA-3’) and reserve (5’-GCCTGTTGTGAGCCTCCTGTCGAA-3’). In the SELEX reaction, 5 mg/mL of purified rPrP^90-231^ fused to an N-terminal six-histidine tag was immobilized in a nickel-loaded HisTrap HP affinity column (GE Healthcare). DNA library was added to the resin, previously equilibrated in 5 mM Tris buffer (pH 7.4) and incubated for 1 h at room temperature. Unbound or low-affinity DNA sequences were removed with low salt buffer [5 mM Tris (pH 7.4), 150 mM NaCl]. Elution of potentially high-affinity sequences was performed in two steps. First, elution was carried out with high (1.5 M) NaCl concentration and then with 1 mL of 3.0 M NaSCN. Salt was removed from the eluted samples with Vivaspin 500 ultrafiltration system (MW *cutoff*: 3000; GE Healthcare), and samples were amplified by PCR using the primers from the DNA library (0.25 mM forward primer and 25 mM reserve primer). Amplified samples were subjected to new rounds of SELEX, with a total of 5 rounds to ensure that resultant sequences exhibited high affinity and specificity for the target. In the last SELEX round, elution was performed with a salt gradient ranging from 1 to 1.5 M NaCl, and samples were collected separately. DNA sequences that eluted at higher salt concentration were further cloned and sequence-verified.

### DNA Cloning and Sequencing

Selected fractions were cloned in chemically-competent *E. coli* Top10 using pcRH2.1TOPOH vector from the TOPOH TA cloning kit (Thermo Fisher Sci., USA). Positive colonies were grown in 25 mL of LB medium at 37 °C under constant shaking. Plasmidial DNAs were isolated with mini-prep kit (Thermo Fisher Sci., USA). DNA sequence analysis was performed by Sanger method using the big dye terminator cycle sequencing kit (Thermo Fisher Sci., USA) on an ABI 3130 Genetic Analyzer (Thermo Fisher Sci., USA). Electropherograms (44 in total, as the fractions were sequenced in both directions, forward and reverse, using T3 and T7 primers present in the cloning vector) were analyzed using BioEdit 7.2.5 and obtained DNA sequences were compared with the reference sequence (primers from the DNA aptamer library: forward 5’-GGGAGACAAGAATAAACGCTCAA-3’ and reverse 5’-GCCTGTTGTGAGCCTCCTGTCGAA-3’). Aptamer sequences were obtained using Jalview 2.8.1 (http://www.jalview.org/) by aligning sequences with flanking primers. Only high-quality electropherograms were selected, in which the forward and reverse pair coincided, even though samples of the same plasmid were sequenced separately. By analyzing the alignments, it was possible to infer that two sequences were complementary, proving the high-quality sequencing for these aptamers. Therefore, these two sequences (named A1 and A2) were selected for synthesis and binding validation.

### Synthetic DNA Oligonucleotides

The single-stranded DNA sequences obtained from SELEX were synthetized and purified by Integrated DNA Technologies (Coralville, USA). DNA oligonucleotides investigated here are listed in **Table 1**. For all experiments, DNA samples were annealed by heating to 95°C for 5 min followed by slow cooling and then kept at room temperature overnight. The concentration of DNA samples was determined by absorbance at 260 nm using the corresponding molar extinction coefficient for the folded oligonucleotide (36).

**Table 1:**
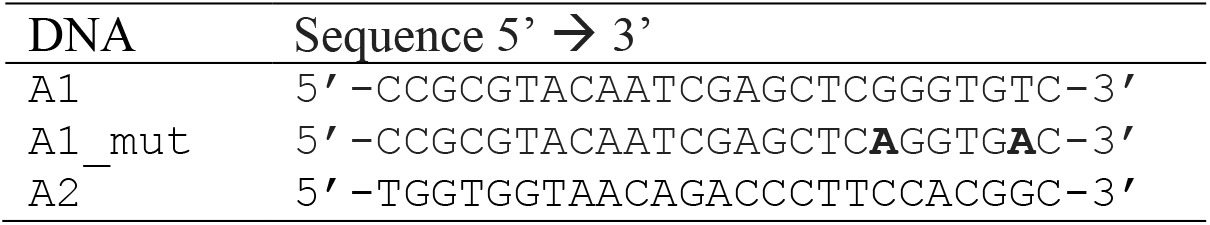
DNA sequences used in the present study.

### UV-Thermal Denaturation

UV-thermal denaturation experiments were performed on a UV-1800 Shimadzu spectrophotometer equipped with a thermostated bath TE-2005. DNA aptamers (A1 at 1.95 μM; A2 at 2.2 μM; A1_mut at 2.6 μM) were dissolved in 20 mM sodium cacodylate (pH 7.0) with or without 100 mM NaCl. For measurements, 1 cm-path length quartz cuvettes with 700 μL volume were used, covered with mineral oil to avoid evaporation. The temperature of the cell holder was increased from 20 to 85 °C at a rate of 0.5 °C/min and absorbance profiles were monitored at 260 nm.

### Isothermal Titration Calorimetry (ITC)

Binding studies were performed at 25 °C on an ITC-200 microcalorimeter (Malvern MicroCal, USA). Both DNA and protein samples were dissolved in 20 mM sodium cacodylate (pH 7.0) with or without 100 mM NaCl. For titrations with A1, A2, and A1_mut, the syringe was filled with DNA solutions at 33, 33 and 100 μM, respectively, while 5 μM rPrP was placed into the sample cell. The reference cell was loaded with degassed ultra-pure water. During titrations, the solution was stirred with a syringe at 750 rpm inside the sample cell without foaming. 1.5 μL aliquots of A1 and A2 aptamers were added sequentially at an interval of 200 s. For A1_mut, a single injection of 10 μL was applied to the cell at an interval of 180 s. The amount of heat generated per titration was determined by integrating the area under the peaks. Heat of dilution of the DNAs in buffer was measured separately and subtracted from the titration data. Analysis of ITC data was carried out using Origin 7.0 provided by MicroCal employing a one-site binding model for curve fitting.

### Nuclear Magnetic Resonance (NMR)

1D ^1^H NMR spectra were recorded on a Bruker Avance III HD 900 MHz spectrometer equipped with an inverse-detection triple resonance z-gradient TXI probe. All experiments were performed at 25°C with 100 μM A1, A2, and A1_mut in 20 mM sodium cacodylate (pH 7.0) with and without 100 mM NaCl, and 5% D_2_O. Data were acquired and processed using TopSpin 3.2 (Bruker Biospin).

### Circular Dichroism (CD)

Secondary structure of A1, A2, A1_mut, and their interaction with rPrP^90-231^ was monitored by CD using a Chirascan spectropolarimeter (Applied Photophysics, Surrey, UK). Spectra were collected at 25°C in a 1 mm-path length quartz cuvette. For free DNA oligonucleotides, CD measurements were performed with 50 μM aptamer in 20 mM sodium cacodylate (pH 7.0) with or without 100 mM NaCl. For protein:DNA interaction studies, 10 μM rPrP^90-231^ in 20 mM sodium cacodylate (pH 7.0) was titrated with increasing concentrations of aptamers (2, 5 and 10 μM). Four scans were collected over the wavelength range of 200 to 340 nm (free aptamers) and from 200 to 260 nm (rPrP^90-231^:aptamer interaction) at a scanning rate of 100 nm/min, 0.1 s per point, and 1 nm bandwidth. All spectra were subtracted from the corresponding buffer spectrum.

### Small Angle X-ray Scattering (SAXS)

SAXS data were collected at the SAXS1 beamline at the Brazilian Synchrotron Light Laboratory (CNPEM, LNLS, Campinas, Brazil). Scattering was recorded with a two-dimensional pixel detector (Pilatus 300K) using a mica sample holder. The sample to detector distance was 1 m, resulting in a scattering vector range of 0.01 < q < 0.45 Å^−1^, where q is the magnitude of the q-vector defined by q = 4π sin (θ)/λ (2θ is the scattering angle). Data were collected at 25 °C with 2 mg/mL A1, A2, and A1_mut in 20 mM sodium cacodylate (pH 7.0). The background buffer scattering was subtracted from the corresponding sample scattering curve. The radius of gyration (R_g_) of the aptamers was obtained from the linear regression of Guinier plots in the appropriated *q* range (*q* < 1.3/R_g_). Globularity and overall shape of the samples was inferred from the Kratky plots (*q*^2^.I(*q*) vs. *q*) (37).

### Differential Interference Contrast (DIC) and Fluorescence Microscopy

Droplet formation was studied by DIC and fluorescence microscopy. Samples of 10 μM rPrP^90-231^ were incubated with 2 to 10 μM of A1, A2, and A1_mut aptamers. To investigate physicochemical characteristics of protein-rich condensates, 10% (w/v) PEG-4000 (Thermo Scientific, USA) or 10% (w/v) 1,6-hexanediol (Merck, Germany) were mixed with 10 μM rPrP^90-231^ in 10 mM Tris (pH 7.4), 100 mM NaCl. Samples were maintained at room temperature 10 min prior to acquisition of micrographs, unless specified. Image chambers for microscopy were assembled by positioning one layer of double-sided tapes at the center of a glass slide. Subsequently, a coverslip was gently pressed on top of the sticky tapes generating a spacer with uniform thickness. Immediately after homogenization, 20 μL of samples were pipetted through the chamber and settle to stand for 10 minutes with coverslip facing down. Micrographs were acquired under Köhler illumination on inverted confocal microscope (Leica TCS SPE, Leica Microsystems, Germany) with a 63 × objective (oil immersion). For DIC, a 488 nm argon laser was used. To investigate amyloid autofluorescence, samples were excited at 405 nm and fluorescence signals were collected at the 450-500 nm range. The presence of solvent-exposed hydrophobic residues was assessed by SYPRO orange staining (5000 × stock, Sigma-Aldrich, USA). SYPRO orange reagent was diluted (1:100) in 44 μM rPrP^90-231^ sample and incubated for 1 h prior to imaging (excitation at 488 nm; emission at the 500-650 nm range). All images were collected at room temperature and processed with Fiji (distribution of the ImageJ software, USA) using histogram stretching to improve contrast. To quantify the number of droplets at each experimental condition, a central area of 0.1 mm^2^ was equally positioned in three fields of view of each sample. Next, the droplets inside this area were manually marked followed by automated identification and counting using Fiji. Droplets diameter was determined using 50 droplets comprised in 5 to 10 images. Data were analysed with GraphPad Prism (version 8.1.1; GraphPad Software).

### Transmission Electron Microscopy (TEM)

All samples were prepared in 10 mM Tris (pH 7.4), 100 mM NaCl. TEM images were performed with 10 μM rPrP (rPrP^90-231^ and rPrP^23-231^) in the presence of 2 μM DNA aptamers (A1, A2, and A1_mut) (5:1 molar ratio). Grids were prepared after incubation at 25 °C at two different times, 0 and 24 h. Samples were adhered to a formvar/carbon coated grid and then stained with 2% uranyl acetate solution. Images were acquired with JEOL JEM1011 electron microscope (Electron Microscopy Platform Rudolf Barth, Instituto Oswaldo Cruz, Rio de Janeiro, Brazil).

### Nanoparticle Tracking Analysis (NTA)

Particle size and concentration were assessed by NTA, carried out with a NanoSight NS300 (Malvern Instruments, Worcestershire, UK) equipped with a sample chamber, a sCMOS camera and a 532 nm laser. The assemblies of rPrP^90-231^ and A1, A2, and A1_mut were observed at a 5:1 protein:aptamer molar ratio, using rPrP^90-231^ and DNAs at a final concentration of 5 and 1 μM, respectively. All samples were prepared in 10 mM Tris (pH 7.4), 100 mM NaCl. At least five readings were collected for each sample at 25 °C.

### Statistics

Analyses were performed in GraphPad Prism (version 8.1.1; GraphPad Software) using a one-way ANOVA Bonferroni multiple comparisons test. Results are expressed as the mean and the error bars indicate standard deviation (SD).

## RESULTS

### High-affinity rPrP^90-231^ ligands selected by SELEX

In previous studies, three different sites of PrP interaction with DNA were identified by our group and others, being: two lysine clusters located in the N-terminal flexible domain, between residues 23 to 27 and 101 to 110 (22–24), and a third binding site in the C-terminal globular domain (residues ~121-231), in which specific residues were not assigned (23, 38). Herein, we searched for aptamers against the C-terminal domain of recombinant murine PrP (amino acids 90-231), which includes part of the N-terminal region containing the second lysine cluster. We performed the selection of aptamers by SELEX, which enabled us to isolate DNA sequences that bind rPrP^90-231^ with high affinity.

Briefly, DNA aptamers were selected against in-column immobilized histidine-tagged rPrP^90-231^ after 5 rounds of the SELEX protocol. Aptamers were eluted with different salt concentrations, desalted and amplified by PCR. Agarose gel electrophoresis revealed that eluted sequences presented the expected mass, according to the aptamer library (~50 bases including the flanking regions) (**Fig. S1**). Aptamers eluted at the highest salt concentration (1.5 M) were cloned into expression vectors and transformed in *E. coli*. Each aptamer-positive colony was grown individually. Plasmids were isolated and inserts were sequenced using primers T3 and T7 from the cloning kit. Only high-quality electropherograms that yielded sequences whose identity was confirmed by complementarity of sense and antisense bases performed in different wells and properly aligned with primers from the aptamer library were selected (**Fig. S2**). Thus, aptamer sequences named A1 and A2 (**Table 1**) were sent for synthesis to further validate high-affinity interaction with rPrP.

### Thermodynamic analysis of rPrP:aptamer interaction

The affinity of the ligands selected by SELEX and the thermodynamic basis of their interaction with rPrP were determined by ITC. Titration of A1 into rPrP^23-231^ and rPrP^90-231^ revealed average dissociation constants *K*_d_ of 838 and 232 nM, respectively (**Fig. 1**, A and B), indicating binding selectivity to rPrP. The similarity in binding affinities for both rPrP constructs suggests that the N-terminal intrinsically disordered region does not significantly contribute to aptamer interaction. Therefore, we can conclude that the C-terminal globular domain represents the high-affinity aptamer-binding region of rPrP. For A2, interaction with full-length rPrP (rPrP^23-231^) yielded a *K*_d_ of 1.47 μM, while interaction with the C-terminal construct rPrP^90-231^ occurred with a *K*_d_ of 1.93 μM (**Fig. 1**, C and D). Binding affinity for A1 is ~10-fold higher than that for A2, suggesting the specific recognition of A1 by the globular domain of rPrP, which may be related to both sequence and conformation of the aptamer. Binding stoichiometries were variable (from 3:1 to 6:1 protein:DNA ratio); high-affinity interaction of rPrP^90-231^ with A1 revealed a 5:1 stoichiometry. **Table 2** summarizes the thermodynamic parameters obtained from the ITC data. Negative enthalpies suggest that the binding process is enthalpically driven.

**Table 2.** Thermodynamic parameters obtained from ITC experiments for rPrP:aptamer binding.

| rPrP:aptamer | N | $K_d (\times 10^{-6} \text{ M})$ | $\Delta H (\text{kcal mol}^{-1})$ | $\Delta S (\text{kcal mol}^{-1} \text{ deg}^{-1})$ |
| --- | --- | --- | --- | --- |
| rPrP<sup>23-231</sup>:A1 | 0.19 | $838 \times 10^{-3}$ | -0.013 | -0.41 |
| rPrP<sup>90-231</sup>:A1 | 0.18 | $232 \times 10^{-3}$ | -0.013 | -0.40 |
| rPrP<sup>23-231</sup>:A2 | 0.35 | 1.47 | $-2.33 \times 10^{-6}$ | $-7.79 \times 10^{-3}$ |
| rPrP<sup>90-231</sup>:A2 | 0.16 | 1.93 | $-0.11 \times 10^{-6}$ | -0.35 |

**Figure 1.**
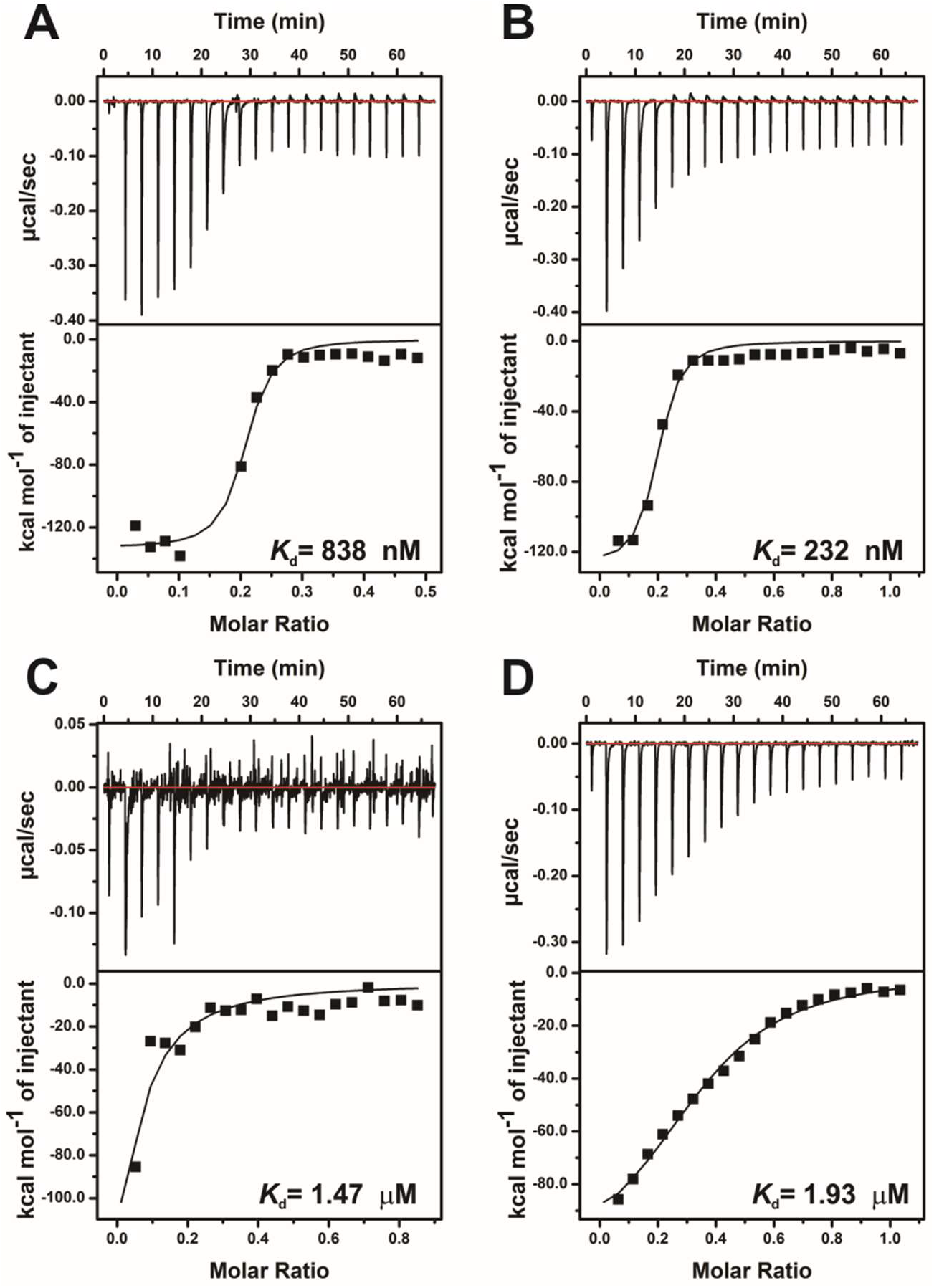
ITC binding isotherm of rPrP^23-231^ and rPrP^90-231^ with selected aptamers. Raw ITC data (top panel) and the integrated heat values plotted against rPrP:aptamer molar ratio (lower panel). (**A**) rPrP^23-231^:A1. (**B**) rPrP^90-231^:A1. (**C**) rPrP^23-231^:A2. (**D**) rPrP^90-231^:A2. Experiment was performed with 5 μM rPrP in 20 mM sodium cacodylate (pH 7.0) at 25 °C.

Protein:DNA interaction is mediated by electrostatic contacts with the phosphate backbone as well as by hydrogen bonds and van der Waals interactions with the specific nitrogenous bases (39). To gain further insights into the specificity of interaction of rPrP^90-231^ with the SELEX-selected DNA aptamers, binding isotherms were acquired at increasing NaCl concentrations (**Fig. S3**). Increasing NaCl concentration led to a decrease in DNA binding affinity, as expected for systems in which electrostatic interactions participate in complex formation. However, despite the reduction in affinity, rPrP^90-231^ was able to bind A1 in the presence of 500 mM NaCl (*K*_d_ 18 μM; **Fig S3B**), while A2 binding, in the same condition, was not maintained (**Fig. S3D**). This difference in salt susceptibility suggests a difference in the mechanism of aptamer recognition by rPrP^90-231^, highlighting a stronger affinity and specificity for A1.

### Structural characterization of A1 and A2 aptamers

To further investigate the molecular mechanisms underlying rPrP^90-231^ interaction with A1 and A2, we conducted a structural characterization of aptamers using different biophysical techniques. CD spectrum of A1 in the absence of salt consisted of two major low-intensity signals, a shortwave negative (243 nm) and a longwave positive (282 nm) band, characteristic of a B-type duplex (**Fig. 2A**). In the B form, the base pairs are nearly perpendicular to the helix axis, resulting in a CD signal with reduced intensity (40). In addition, 1D ^1^H NMR spectrum of A1 revealed the presence of non-exchanging, Watson-Crick hydrogen bonded imino resonances, indicating intrastrand base pairing and the formation of a hairpin structure (**Fig. 2B**). Addition of 100 mM NaCl changed the CD spectrum of A1, leading to an intensity increase in the shortwave negative band (242 nm) and the appearance of two low-intensity longwave positive bands (261 and 288 nm), suggesting a transition from the canonical B to a nonusual B’ heteronomous structure (**Fig. 2A**). This form is characterized by a more unwound double helix and, consequently, a narrower minor groove (41). Previous reports have shown that low salt concentration favors DNA conformations exhibiting a narrower minor groove, owing to the greater phosphate repulsion (40, 42). In the presence of salt, Watson-Crick imino resonances were maintained (**Fig. 2B**). CD spectrum of A2, regardless of the addition of salt, was characterized by signals typical of a B-type helical structure: two high-intensity bands, one negative (243 nm) and one positive (277 nm) (**Fig. 2C**). 1D ^1^H NMR spectrum of A2 showed no evidence for paired imines (**Fig. 2D**), suggesting a single-stranded coil conformation with no secondary structure. Kratky plots from SAXS data showed that A1 adopts a globular structure, while A2 is unfolded (**Fig. 2E**) (37, 43). Through the Guinier plot, we estimated a radius of gyration of 1.77 nm for A1 (**Fig. S4**). The hydrodynamic radii of both aptamers were calculated using DLS, corresponding to 2.7 nm for A1 and 4.2 nm for A2. Moreover, A1 showed a sigmoidal thermal denaturation curve, enabling us to calculate a Tm of 45 °C, independent of the addition of salt (**Fig. S5A** and **S5D**). For A2, a Tm value could not be measured within the temperature range used in the experiment (20-85 °C) (**Fig. S5B** and **S5E**), suggesting a greater stability for the hairpin-structured A1 aptamer. Our results agree well with the prediction of secondary structure for A1, which indicated that the lowest-free-energy conformation, −7.80 kcal/mol, is structured as a hairpin formed by a hydrogen bond-stabilized stem and an internal loop containing 12 nucleotides (**Fig. 2F**). In contrast, for A2, the calculated minimum Gibbs free energy for a hairpin structure was −1.70 kcal/mol (44).

**Figure 2.**
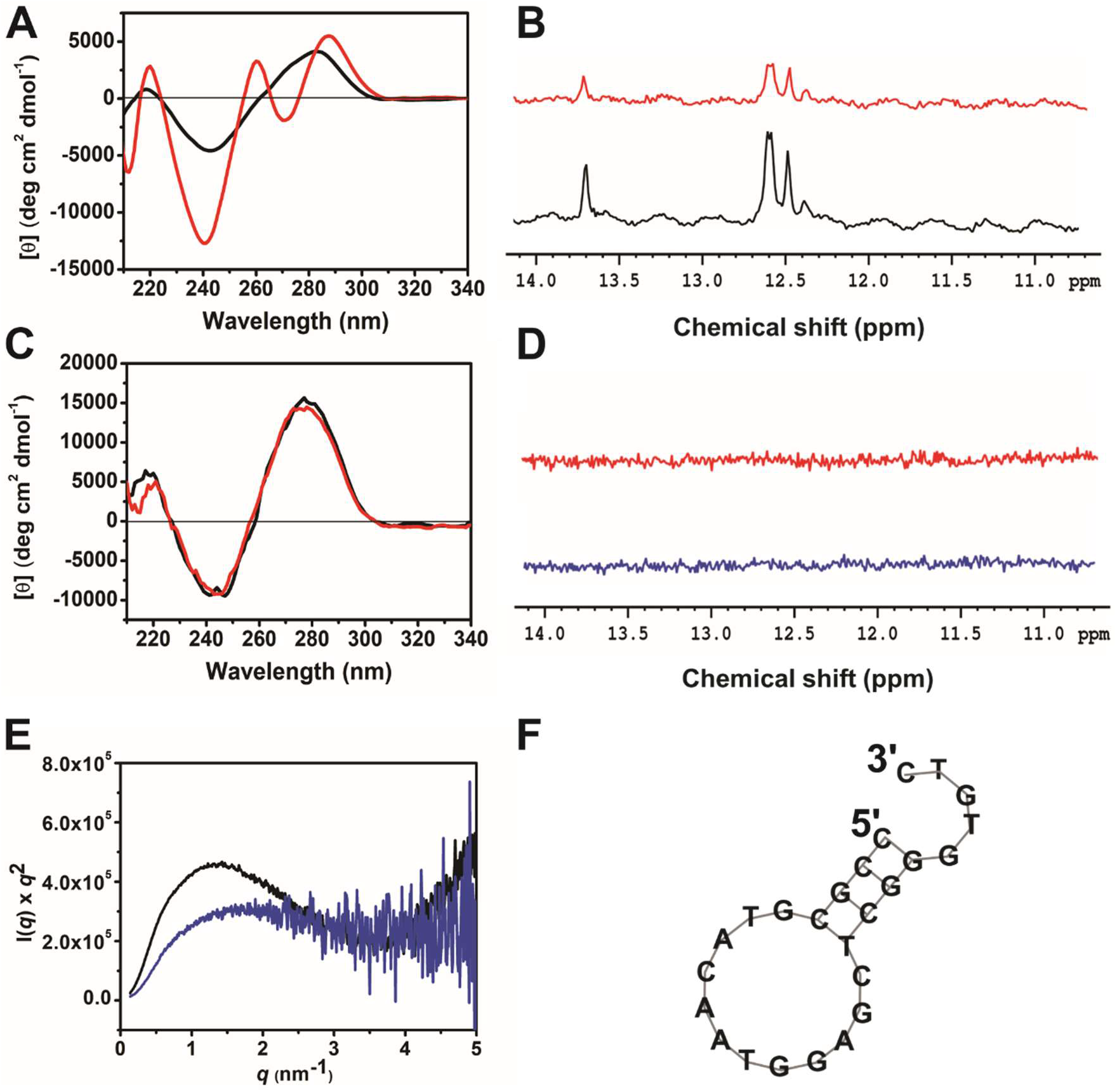
Structural characterization of A1 and A2 aptamers. (**A**) Circular dichroism spectra of 50 μM A1 in 20 mM cacodylate (pH 7.0) (black) and 50 μM A1 in 20 mM cacodylate (pH 7.0), 100 mM NaCl (red), 25 °C. (**B**) 1D ^1^H NMR spectra displaying the chemical shift region of paired imines. Experiment was performed with 100 μM A1 in 20 mM phosphate (pH 7.0), 25 °C. A1 in the absence (black) and presence of 100 mM NaCl (red). (**C**) Circular dichroism spectra of 50 μM A2 in 20 mM cacodylate (pH 7.0) (blue) and 50 μM A1 in 20 mM cacodylate (pH 7.0), 100 mM NaCl (red), 25 °C. (**D**) 1D ^1^H NMR spectra displaying absence of chemical shifts characteristic of paired imines. Experiment was performed with 100 μM A2 in 20 mM phosphate (pH 7.0), 25 °C. A2 in the absence (blue) and presence of 100 mM NaCl (red). (**E**) Kratky plot of A1 (black) and A2 (blue) at 200 μM in 20 mM sodium cacodylate (pH 7.0). (**F**) Prediction of secondary structure for A1, using the Vienna RNAfold package.

### High-affinity interaction is dependent on aptamer structure

To pinpoint the role played by aptamer structure in high-affinity binding to rPrP^90-231^, we made use of an A1 mutant (A1_mut) in which two nucleotides were substituted with the aim of disrupting the base pairing; guanine at position 19 (G19) and thymine at position 24 (T24) were replaced by adenines (**Table 1**). The structural properties of A1_mut were evaluated by various biophysical techniques. CD spectrum of A1_mut was highly similar to that of A2, characterized by a shortwave negative (248 nm) and a longwave positive (277 nm) band of high intensity (**Fig. 3A**). 1D ^1^H NMR spectrum of A1_mut, in the absence of salt, showed no evidence for Watson-Crick hydrogen bonded imines, indicating a single stranded coil conformation. Addition of 100 mM NaCl led to the appearance of resonance signals with very small relative intensity in the region of paired imines (**Fig. 3B**), suggesting a coil-hairpin equilibrium. However, the CD spectrum in this condition is nearly identical to that of A1_mut in the absence of salt, suggesting that the equilibrium is shifted toward the coil conformation. The Kratky plot from SAXS data for A1_mut suggested a less folded structure when compared to A1 (**Fig. 3C**). Moreover, the stability of A1_mut was also investigated by thermal denaturation (**Fig. S5C** and **S5F**). Like A2, the thermal denaturation curve of A1_mut was mostly linear, hindering the measurement of a Tm within the temperatures tested (20-85 °C), regardless of the addition of salt. This is another evidence that A1_mut adopts no regular secondary structure. Next, rPrP^90-231^:A1_mut interaction was investigated by ITC. Despite numerous efforts, we were not able to obtain a reliable binding isotherm with multiple injections for A1_mut at the same conditions used for A1 and A2, most probably due to lower affinity toward rPrP^90-231^. To overcome this problem, protein concentration was increased; however, this resulted in protein aggregation inside the ITC cell. Thus, a single injection of A1_mut into rPrP^90-231^ solution was performed indicating interaction (**Fig. S6**).

**Figure 3.**
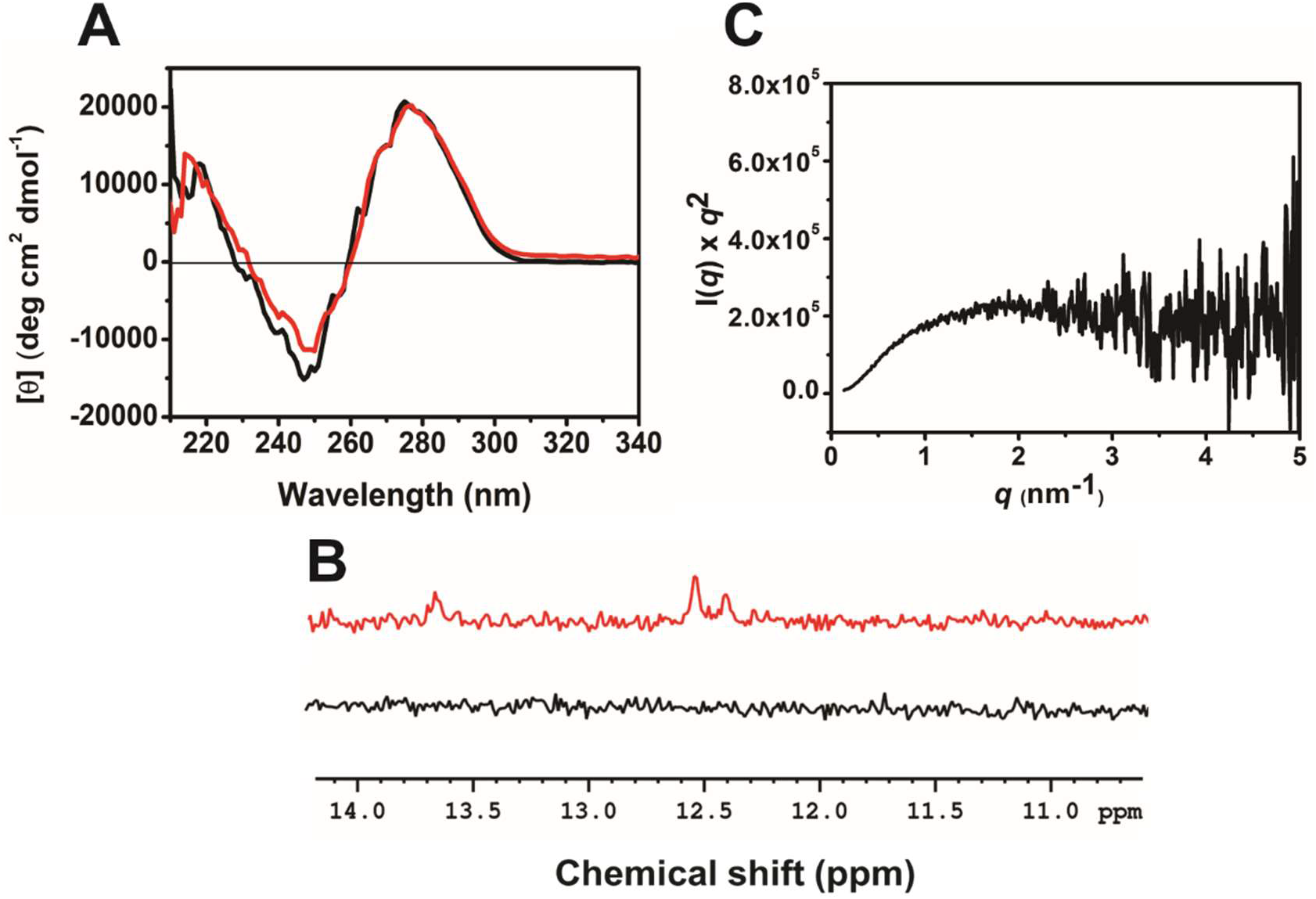
A mutant form of A1 (A1_mut) adopts no ordered secondary structure. (A) Circular dichroism spectra of 100 μM A1_mut in 20 mM cacodylate (pH 7.0) (black) and 20 mM cacodylate (pH 7.0), 100 mM NaCl (red), 25 °C. (B) 1D ^1^H NMR spectra displaying the chemical shift region of paired imines. Experiment was performed with 100 μM A1_mut in 20 mM phosphate (pH 7.0), 25 °C. A1_mut in the absence (black) and presence (red) of 100 mM NaCl. (C) Kratky plot of A1_mut at 200 μM in 20 mM sodium cacodylate (pH 7.0).

### Aptamer binding triggers rPrP^90-231^ loss of secondary structure

To investigate the effect of aptamer binding on rPrP structure, CD experiments were performed at increasing rPrP^90-231^:DNA molar ratios. To guarantee that only the protein signal was observed, we subtracted each CD spectrum from the correspondent free DNA spectrum obtained at the specific concentration. Titration of A1 into rPrP^90-231^ induced a loss of CD signal in a stoichiometry-dependent manner, either caused by protein aggregation and/or partial unfolding. Strikingly, this secondary structure loss was inversely correlated with DNA concentration, so that low DNA concentration (5:1 rPrP^90-231^:DNA molar ratio) resulted in greater signal loss (**Fig. 4A**). Similar results were obtained for A2 (**Fig. 4B**). In contrast, addition of A1_mut did not cause comparable changes to the protein secondary structure at the different stoichiometric ratios evaluated (**Fig. 4C**). To further investigate the molecular events underlying this loss in CD signal, 2D [^1^H, ^15^N] HSQC spectra of PrP^90-231^ were collected upon addition of A1 aptamer (**Fig. S7**). Even in the lowest NA:protein molar ratio (1:10), addition of A1 led to disappearance of most of the ^1^H,^15^N resonance signals, suggesting that: (*i*) A1 binding leads to rPrP^90-231^ aggregation; (*ii*) A1 binding leads to a collection of partially unfolded rPrP^90-231^ states that interconvert in the intermediate NMR time scale. At present, we cannot rule out any of the two mechanisms or even a combination of both; however, the absence of resonance signals at the 2D [^1^H, ^15^N] HSQC spectrum of PrP^90-231^ in the presence of A1 (**Fig. S7**) suggests that the lack of secondary structure observed in the CD spectra is not due solely to protein unfolding.

**Figure 4.**
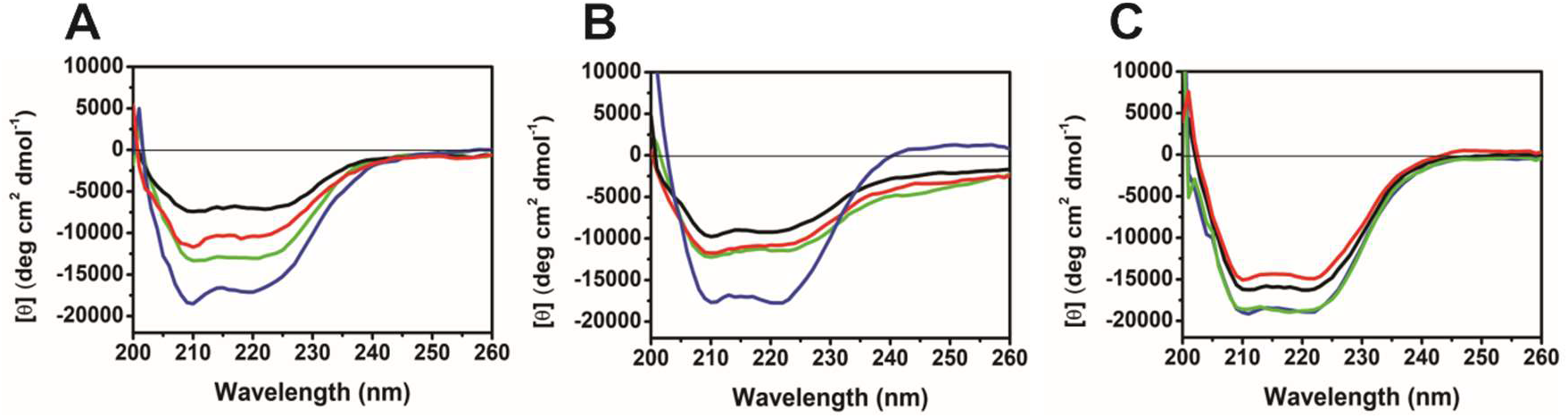
A1 and A2 decrease the characteristic secondary structure signals of rPrP^90-231^. Circular dichroism spectra of rPrP^90-231^ (blue), and at 1:1 (green), 2:1 (red) and 5:1 (black) rPrP^90-231^:DNA molar ratios. (**A**) rPrP^90-231^:A1. (**B**) rPrP^90-231^:A2. (**C**) rPrP^90-231^:A1_mut. Experiments were performed with 10 μM rPrP^90-231^ in 20 mM sodium cacodylate (pH 7.0) at 25°C.

Nanoparticle tracking analysis (NTA) identified distinct particle size and concentration for rPrP^90-231^ bound to the different aptamers. For instance, rPrP^90-231^:A1 complex displayed a larger population of increased size (50-380 nm) than that observed for rPrP^90-231^:A2 and rPrP^90-231^:A1_mut complexes. Interestingly, rPrP^90-231^ binding to A2 or A1_mut yielded nanoparticles distributed within 10-50 nm (**Fig. S8**). Total particle concentration was significantly higher for rPrP^90-231^:A1 than for rPrP^90-231^:A2 or rPrP^90-231^:A1_mut complexes, which showed similar concentrations (**Fig. S8, inset**). The observed size distribution indicate the presence of large macromolecular assemblies, not compatible with 1:1 or 5:1 rPrP:aptamer complexes.

During the attempts to obtain concentrated and soluble rPrP:aptamer complexes, we verified turbidity immediately upon aptamer addition to rPrP samples (**Fig. S9**). Intriguingly, incubation at room temperature for ~7 h led to sample disaggregation. Similar results were observed for full-length rPrP in the presence of a high-affinity RNA sequence (19) and for Syrian hamster rPrP^90-231^ in the presence of an 18-mer DNA sequence (23).

### The C-terminal globular domain of rPrP phase separates *in vitro*

Based on the dynamic aggregation properties of the rPrP:DNA complexes, specifically at the 5:1 rPrP:aptamer molar ratio, we next argued whether liquid-liquid phase separation contributes to the observed effects. Many characteristics of proteins that undergo LLPS is indeed common to PrP: prion protein is an IDP, binds NAs, and possesses low-complexity domains (16). rPrP^90-231^ is able to form multivalent interactions with NA and contains specific polar residues interleaved by aromatic amino acids that can mediate phase separation (**Fig. S10A**) (45). Additionally, rPrP^90-231^ exhibits other typical phase separation-prone characteristics, such as predicted intrinsic disorder (**Fig. S10B**), strong propensity to undergo LLPS (catGranule score 1.946) (**Fig. S10C**), weak alternating charged regions (**Fig. S10D**).

DIC microscopy showed that rPrP^90-231^ phase separates *in vitro* into micrometer-sized spherical condensates with varying diameters in water and buffer (10 mM Tris (pH 7.4), 100 mM NaCl) (**Fig. 5A and 5B**). Phase separation occurred even in the absence of macromolecular crowding agents, albeit 10% PEG 4000 increased droplets diameter compared to rPrP^90-231^ in water (**Fig. 5B**), probably aiding fusion of condensates. Larger rPrP^90-231^ condensates were observed reaching approximately 100 μm of diameter (**Fig. S11A**). Coalescence into large droplets is thermodynamically driven since the net surface tension is diminished over small condensates, *i.e.* Ostwald ripening (46). However, the number of droplets was significantly decreased in both buffer and in the presence of PEG compared to water, suggesting dependence of ionic strength on phase separation behavior (**Fig. 5B**). We were able to follow fusion and fission events characteristic of protein liquid droplets (**Fig. S11B and movie S1**). Additionally, incubation of rPrP^90-231^ with 10% 1,6-hexanediol, an aliphatic alcohol known to disrupt weak hydrophobic contacts involved in LLPS (47), completely abolished droplet formation, confirming phase separation propensity of rPrP^90-231^ (**Fig. S11B**). To provide insights into the structure of the rPrP^90-231^ phase-separated state, we added SYPRO orange, a dye that only exhibits fluorescence when bound to hydrophobic surfaces exposed to solvent (48). After 10 min incubation, co-partitioning of SYPRO orange within rPrP^90-231^ liquid droplets led to discrete fluorescence (**Fig. 5C**). A marked enhancement of SYPRO orange fluorescence was observed after 60 min incubation (**Fig. 5D**). Firstly, dye trapping by higher droplet viscosity may result in increased fluorescence (49). Secondly, one-hour incubation could evolve transition to a more unfolded state with more exposed sites for SYPRO orange binding. Nonetheless, fluorescence in rPrP^90-231^ protein-rich droplets suggests partial unfolding of the protein in the phase-separated state. Interestingly, these large condensates harbor smaller droplets inside, resembling multi-vesicular bodies (50). Such vacuolized condensates of varying densities are formed in response to system perturbation agents (51). We suggest that 1% (v/v) dimethyl sulfoxide (DMSO), which is the SYPRO orange vehicle, can result in this non-equilibrium state depicted by vacuolized droplets possessing varying densities. In addition, SYPRO orange randomly binds to hydrophobic regions that might be among the interactions driving phase transition.

**Figure 5.**
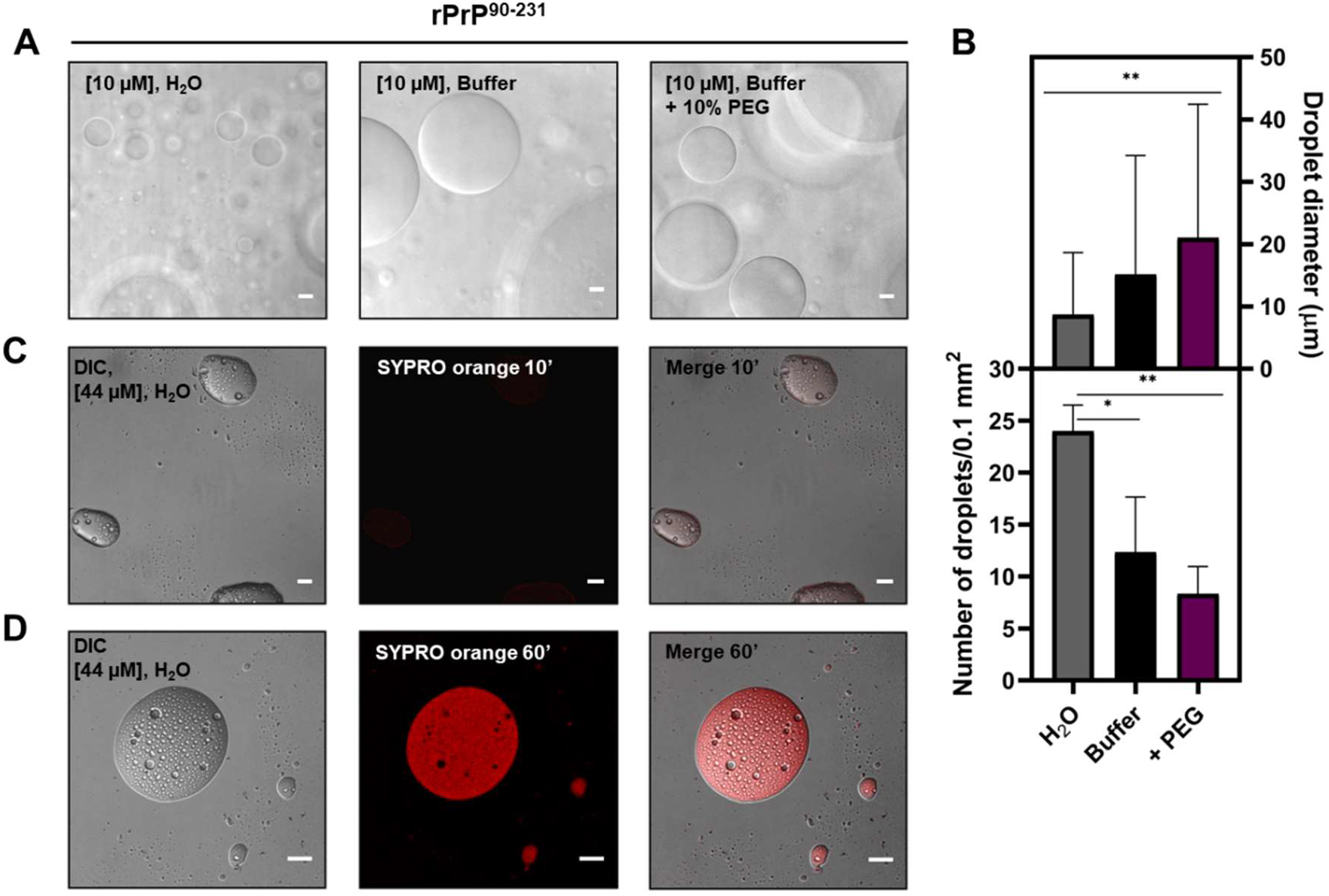
rPrP^90-231^ undergoes liquid-liquid phase separation *in vitro*. Representative images from DIC microscopy of (A) 10 μM rPrP^90-231^ in water (top left), 10 μM rPrP^90-231^ in 10 mM Tris (pH 7.4), 100 mM NaCl (top middle) and 10 μM rPrP^90-231^ in buffer upon addition of 10% (w/v) PEG 4000 (top right), shading depicts floating liquid droplets in other focal planes. All samples were incubated for 10 minutes prior to image acquisition. (B) rPrP^90-231^ droplets quantification from DIC micrograph analyses. rPrP^90-231^ at 10 μM in water (gray column), in buffer (black column) and in buffer containing 10% (w/v) PEG 4000 (purple column). Measurement of liquid droplets diameter (top). Data are represented as the mean ± SD (n=50 droplets). Quantification of rPrP^90-231^ droplets number per 100 μm^2^ area (bottom). Data are represented as the mean ± SD (n=3 images). *p < 0.05, **p < 0.005. (C) and (D) 44 μM rPrP^90-231^ fluorescently stained with SYPRO orange for 10 minutes (C) and for 60 min (D) prior to imaging. Scale bar, 10 μm.

### Aptamers modulate rPrP^90-231^ liquid-liquid phase separation

To explore if the 25-mer oligonucleotides influence rPrP^90-231^ phase separation, we performed phase transition assays in the presence of aptamers. Addition of A1 or A2 to rPrP^90-231^ resulted in a decrease in droplets diameter (15.12 ± 19.10 versus 3.93 ± 1.43 for A1 and 7.20 ± 3.30 for A2, at 5:1 rPrP^90-231^:aptamer molar ratio) and increase in the number of significantly small condensates in a concentration-dependent manner (**Fig. 6A, 6B** and **6D**). Specifically, at 5:1 rPrP^90-231^:aptamer stoichiometry the number of droplets/0.1 mm^2^ was 98.33 ± 9.46 for A1 and 73.66 ± 14.97 for A2, respectively, versus 12.33 ± 5.31 for rPrP^90-231^ in buffer without aptamers (**Fig. 6D**). rPrP^90-231^ liquid droplets were more abundant at lower rPrP^90-231^:A1 or A2 molar ratios (**Fig. 6D**). It was previously shown that LLPS is highly dependent on the protein:NA molar ratio; lower ratios of specific nucleic acids promote phase separation (31). Thus, our data are in accordance with other NA-binding proteins that suffer LLPS. Next, we investigated the effect of A1_mut on rPrP^90-231^ phase transition. Only one liquid droplet was observed in the entire cover of the glass slide (**Fig. 6C**), indicating that this aptamer negatively regulated liquid droplets formation. Decreasing A1_mut concentration by half (2:1 rPrP^90-231^:aptamer molar ratio), resulted in few irregular elongated structures consistent to a less dynamic gel-like state (**Fig. 6C**, black arrows). These condensates were deposited on the glass slide indicating higher viscosity in contrast to A1 or A2:rPrP^90-231^ condensates, which were floating. This abnormal morphology was associated to a different physical state that evolved to solid aggregates over time, i.e., suffering liquid-to-solid transition (30). Structures similar to aggregates were visualized at 5:1 rPrP^90-231^:A1_mut ratio. Isolated aptamers or a non-related protein, BSA, did not phase separate at the same conditions used for rPrP^90-231^ (**Fig. S12**).

**Figure 6.**
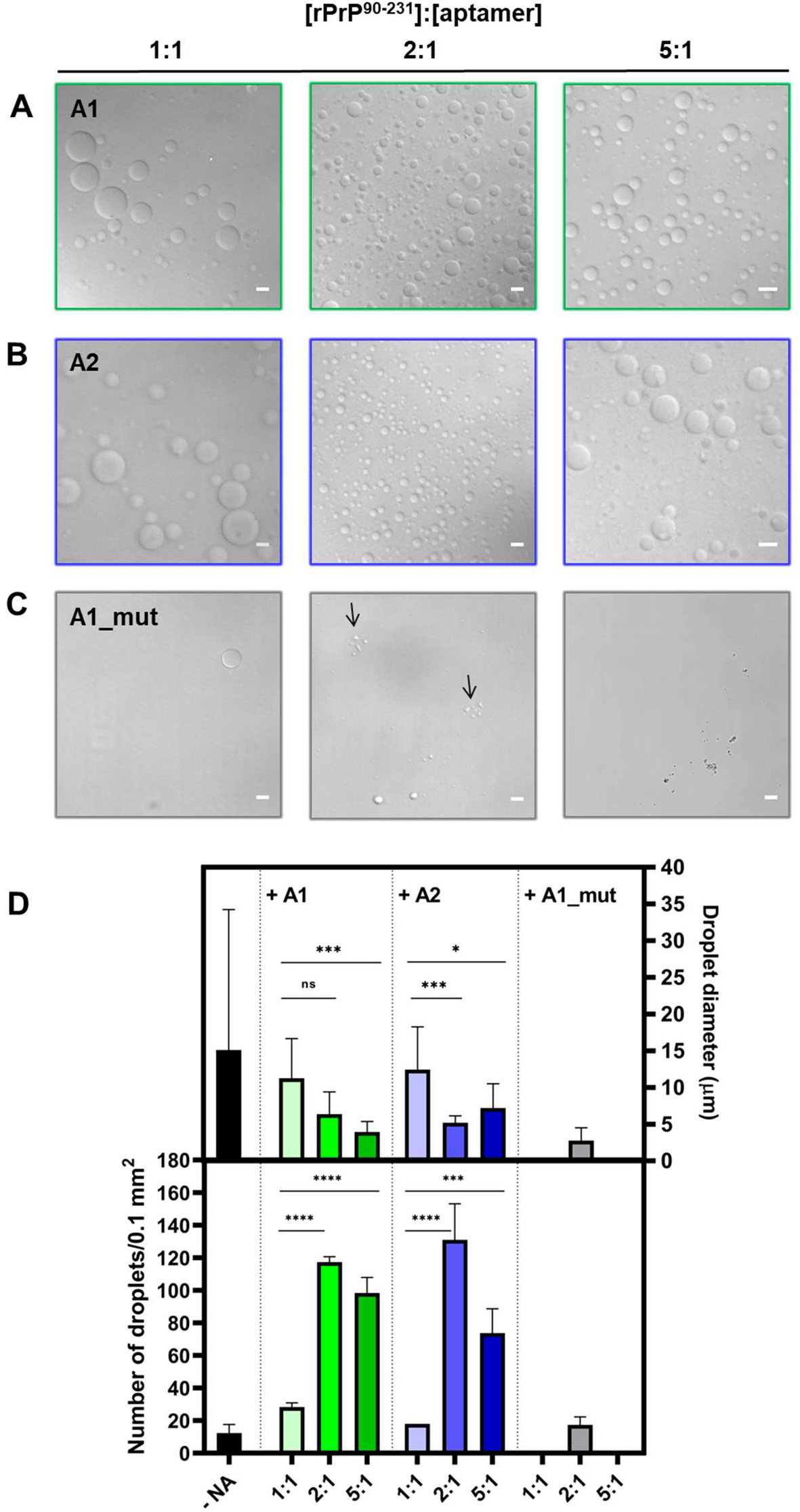
Aptamers tune rPrP^90-231^ phase separation. Representative DIC micrographs of rPrP^90-231^:aptamers after 10 minutes incubation at 1:1, 2:1 and 5:1 molar ratio, respectively, in 10 mM Tris (pH 7.4), 100 mM NaCl in presence of A1 (A), in presence of A2 (B) and in presence of A1_mut (C). (D) Biomolecular condensates composed by rPrP^90-231^ were quantified from DIC images analyses. 10 μM rPrP^90-231^ in buffer without nucleic acids (black column), 10 μM rPrP^90-231^ with addition of increasing concentrations of A1 (shades of green) or A2 (shades of blue) or A1_mut (gray column). Measurement of liquid droplets diameter (top). Data are represented as the mean ± SD (n=50 droplets). Quantification of rPrP^90-231^ droplets number per 100 μm^2^ area (bottom). Data are represented as the mean ± SD (n=3 images). Liquid droplets diameter in the presence of A1 and A2 aptamers. One-way ANOVA/Bonferroni analysis showed significantly results upon incubation of aptamers, *p < 0.05, ***p < 0.001, ****p < 0.0001. Scale bar, 10 μm.

### Aged rPrP^90-231^:A1_mut droplets evolve to amyloid-like fibrillary aggregates

It is well established that aged liquid droplets can mature to fibrillar aggregates *in vitro* (33); thus, we asked whether rPrP^90-231^ liquid condensates transit to solid-like material over time. Within the period of 20 min, we observed tiny bright dots on the surface of rPrP^90-231^ droplets in 10 mM Tris (pH 7.4), 100 mM NaCl. After 60 min, filamentous-like “substructures” were evident on the surface of large rPrP^90-231^ condensates in water (**Fig. S13A**), hence, droplets biophysical properties may change over time. After 6 hours, the liquid droplets assumed characteristics of “aged” droplets, such as darkly refractile appearance and the presence of aggregates sticking out of the condensate (**Fig S13B**). We sought to monitor rPrP^90-231^ aged droplets transition into aggregates in real time. Consistently, aged droplets deform into solid fiber-like structures (**Movie S2**).

It has been shown that a mixture of DNA can induce rPrP^121-231^ unfolding and aggregation into amyloid-like fibers (38). To determine whether A1_mut, which is unstructured according to our biophysical data, promotes formation of rPrP^90-231^ amyloid-like fibers, we explored the characteristic intrinsic fluorescence of amyloids. This approach enables visualization of protein amyloid structures upon excitation in the UV range in a dye-free environment without interference of extrinsic label binding (52). Upon 60 min of A1_mut addition to rPrP^90-231^ (1:1 molar ratio), amyloid structures colocalized with aged droplets (**Fig 7A**). Moreover, at longer time intervals (24 h) the condensates were no longer present, leaving only amyloid fibril-like structures (**Fig 7B and 7C**). Most likely, rPrP^90-231^ in the absence of NA, majorly forms non-amyloid aggregates over time as no auto-amyloid fluorescence was detected (**Fig. S13A**). Conversely, we identified rPrP^90-231^:A1_mut (5:1 molar ration) gel-like elongated droplets at 10 min incubation (**Fig. S13C**, pointed by black arrows) that matured into amyloid-like fibrils after 1 h (**Fig. S14A**). Additionally, intrinsic fluorescence of amyloid fibril-like structures colocalized with SYPRO orange staining, confirming the proteinaceous nature of these structures (**Fig. S14A**). We used α-synuclein fibers as a positive control (**Fig. S14B**). Phase separation of NA-binding proteins has been demonstrated to be dependent upon specific contacts between high affinity NAs that usually contain complex secondary structure (31, 53). Relevantly, A1 and A2 kept rPrP^90-231^ condensates in a dynamic state, preventing transition to solid-like structures in the timeframe examined. In contrast, binding of A1_mut prompted transition of liquid droplets to lower energy amyloid species that mimic pathological solid aggregates. Together, these data indicate that rPrP^90-231^ phase separation is tunable by specific aptamer binding and is dependent on its secondary structure architecture.

**Figure 7.**
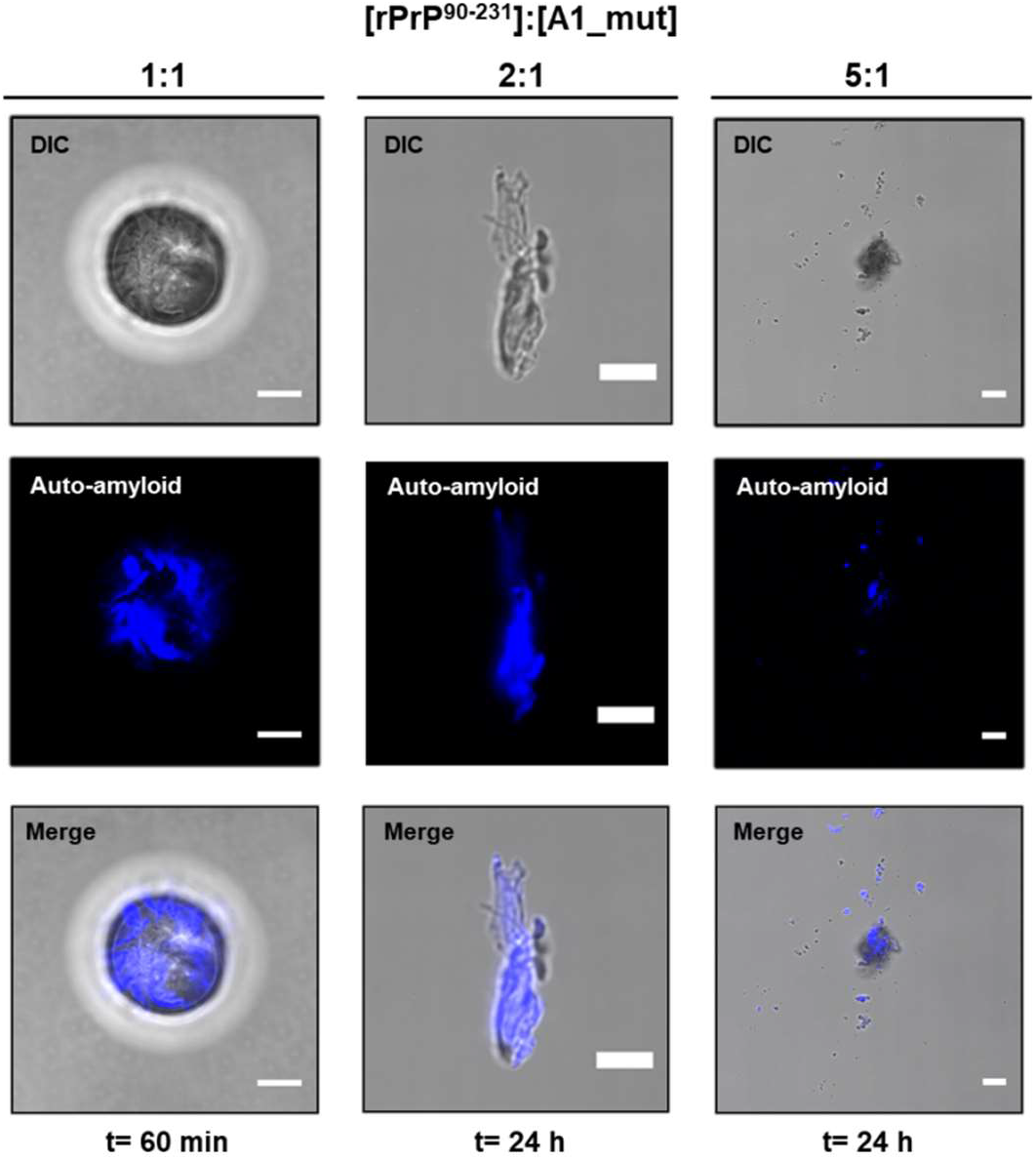
A1_mut promotes transition from rPrP^90-231^ liquid droplets to auto-fluorescent species suggestive of amyloids. Representative DIC and fluorescence micrographs of rPrP^90-231^:A1_mut at 1:1, 2:1 and 5:1, respectively, in 10 mM Tris (pH 7.4), 100 mM NaCl. Left column: 1:1 stoichiometry, 1 h after incubation with A1_mut; middle: 2:1 rPrP^90-231^:A1_mut molar ratio, 24 h after incubation; right column: 5:1 rPrP^90-231^:A1_mut molar ratio, 24 h after mixing. Scale bar, 10 μm.

### Morphology of rPrP^90-231^ condensates depends on aptamer sequence and structure

Circular structures, characterized as droplets, were previously observed by electron microscopy for liquid-liquid phase separated tau protein (48). Thus, after observation of the LLPS phenomenon, TEM was performed to assess the morphological characteristics of the rPrP^90-231^:aptamer condensates and their changes over time. Although TEM micrographs showed that immediately (t=0h) after aptamer addition to rPrP^90-231^ (5:1 rPrP^90-231^:aptamer molar ratio) liquid droplets are formed, the condensate morphology is different depending on the aptamer (**Fig. 8A, C** and **E**). The droplets observed at t=0h after A1 binding to rPrP^90-231^ displayed a more organized structure, with a well-defined electro dense border and an electro lucent central zone. Small aggregates were also seen at the edges of the condensate (**Fig. 8A**). The droplets formed by rPrP^90-231^ binding to A2 and A1_mut were morphologically similar, showing an electron dense center and the absence of a well-defined border as observed for A1 (**Fig. 8C** and **E**). In contrast, after 24 h of incubation, rPrP^90-231^ droplets, at all tested conditions, transitioned to large irregular and branched aggregates with similar morphology regardless of the DNA aptamer (**Fig. 8B, D** and **F**).

**Figure 8.**
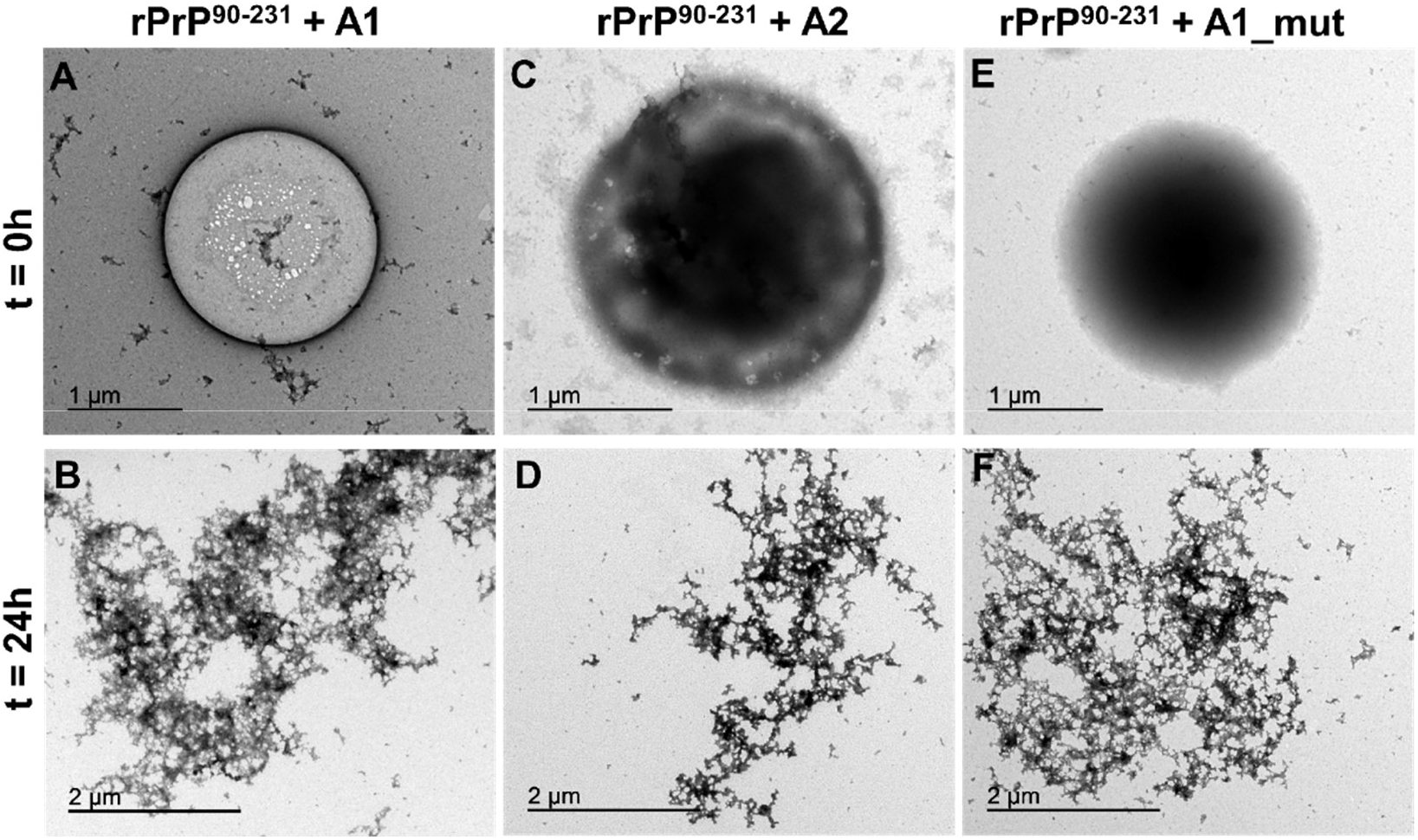
TEM shows that rPrP^90-231^:aptamer condensates evolve to amorphous aggregates over time. Liquid droplets formed by incubation of rPrP^90-231^ in the presence of A1, A2 and A1_mut (5:1 molar ratio) at t=0h. After 24 h incubation, droplets transits to aggregates. Experiments were performed with 10 μM rPrP^90-231^ in 10 mM Tris (pH 7.4), 100 mM NaCl at 25 °C.

## DISCUSSION

Pioneering work has shown that membrane-less organelles (MLOs) assemble via liquid phase separation (54, 55). Importantly, the drivers of LLPS are intrinsically disordered proteins, most of them are directly associated to the pathogenesis of degenerative diseases (56). There is mounting evidence that misregulated phase transition can lead to aggregation-prone protein species (33, 34, 49, 57). PrP possesses most of the physico-chemical characteristics necessary to drive LLPS (**Fig. S10**), including interaction with NAs (16). In addition, PrP-induced ribonucleoprotein granules harbor the characteristics of MLOs (58), pointing toward an essential investigation of PrP demixing properties. NAs can trigger conformational changes in PrP and induce protein aggregation depending on NA structure, size and sequence (14, 21). Moreover, the observed effects on PrP aggregation are highly dependent on the PrP:NA binding stoichiometry (14) (Kovachev *et al.*, submitted). Full-length PrP (PrP^23-231^) contains at least 3 sequence motifs that bind NAs (17), being able to form an interconnected polymer network with NAs. The PrP construct used here, PrP^90-231^, encompasses the second lysine cluster that can interact with NAs, which, when neutralized, results in formation of PrP^Sc^-like aggregates (59). It was hypothesized whether binding to polyanionic cofactors, such as NAs, would lead to charge neutralization of the lysine cluster, promoting formation of scrapie PrP (59).

In accordance to the results shown here, other proteins related to degenerative diseases suffer LLPS induced by NA binding in a clear stoichiometry-dependent fashion (31, 60). High protein:NA ratios, analogous to the cytosol environment, induce larger LLPS effect, whereas low protein:NA ratios, as the nuclei *millieu*, maintain proteins in a dynamic soluble state (31). A recent work described LLPS for full-length PrP^C^ in physiological buffer and that addition of Amyloid-β oligomers resulted in hydrogel formation containing both Amyloid-β oligomers and PrP^C^ (32). Based on this observation and on previous work showing NA-induced PrP aggregation, we hypothesized that NAs can modulate the oligomerization process, nucleating higher order assemblies resulting in biomolecular condensates.

We employed the SELEX methodology to identify high-affinity NA ligands of rPrP^90-231^ and further biophysically characterize the rPrP^90-231^:NA complex. Next, we investigated the interaction of two SELEX-selected, 25-mer aptameric DNA sequences, A1 and A2, and used a mutant form of A1 (A1_mut) to infer whether the NA structure played a role in rPrP binding and resultant conformational changes. Thermodynamic analysis of rPrP:aptamer interaction revealed high-affinity binding. Both A1 and A2 induced loss of protein CD signal, suggesting a combination of molecular events, which may include rPrP^90-231^ aggregation and partial unfolding. Since incubation with A1 aptamer led to clouding (**Fig. S9**) instead of protein precipitation, we evaluated whether rPrP^90-231^ could suffer LLPS and the ability of the SELEX-identified aptamers to modulate this effect. rPrP^90-231^ phase separates alone in aqueous solution (**Fig. 5**) and aptamer binding induces formation of smaller and more abundant liquid droplets (**Fig. 6**). In contrast, incubation of rPrP^90-231^ with the mutated form of A1 that does not adopt a hairpin structure, A1_mut, did not reveal the same extension of LLPS as seen for A1 and A2 (**Fig. 6**). It has been reported that specific NAs containing secondary structure can nucleate formation of biomolecular condensates (53) even in high concentration of background total NA (31). In contrast to highly structured NAs, A1_mut might not provide a scaffold for rPrP^90-231^ LLPS. In fact, rPrP^90-231^:A1_mut condensates at 5:1 molar ratio were organized in solid deposits with characteristic amyloid intrinsic fluorescence (**Fig. 7**). We speculate that A1_mut interacts with rPrP^90-231^ in a non-specific manner, leading to aberrant phase transition that evolves to amyloid aggregates over time.

PrP is mostly found at the cell surface, covalently attached to the membrane by a GPI-anchor (11). Thus, it is debatable how and in which biological compartment PrP^C^ would interact with NAs and undergo LLPS. Nonetheless, micrometer-sized clusters of proteins can occur at membranes and have been associated to synaptic activity (61, 62), addressing a possible role for PrP^C^ phase separation as signaling hubs at the plasma membrane. However, PrP^C^ is not solely found at the extracellular membrane, there is compelling evidence showing PrP alternative subcellular localization, including the cytosol (63, 64). For GPI-anchored proteins, it is well-established that they cycle along the secretory pathway, being translocated to the ER after synthesis, and transported to the Golgi apparatus before being directed to the plasma membrane (64). In specific conditions, PrP was also found in the nucleus (65, 66), highlighting the possibility of a PrP:NA encounter. Furthermore, formation of ribonucleoprotein particles induced by a cytosolic form of PrP was previously described (67). These PrP-induced ribonucleoprotein granules harbor the characteristics of intracellular membrane-less organelles (58), supporting the idea that PrP may suffer LLPS *in vivo* and its misregulation might evolve to pathogenic states. Phase transition driven by the DNA aptamers was dependent on the complex stoichiometry, a characteristic of proteins that suffer LLPS, such as FUS and TDP-43 (31). Accordingly, patterned localization of NAs inside the cellular *milieu* can provide spatiotemporal tuning of PrP phase separation.

Disease-associated aging can lead to loss of efficient nucleus-cytosol shuttling, increasing time of protein residence in the cytosol, where the NA to protein ratio is low(31). Thus, phase separation is favored and this highly dynamic state may constitute a primary event for maturation into aggregates upon stress triggers, similar to TDP-43 demixing (68). Likewise, promising research on PrP^C^ stabilization with a small molecule displaying low micromolar range affinity (69) supports the idea that the high-affinity aptamers described here could compete with cellular stress triggers. Thus, our work provides new insights and paves the way for the rational development of anti-aggregation therapies.

## AUTHOR CONTRIBUTIONS

ASP and YC designed the research; COM was involved in all aspects of the experimental design, data collection, analysis and interpretation. YMP and MJA performed and analyzed all microscopy experiments. BM, MT and SM did the SELEX experiments and analyzed the resulting sequences. OCLB and MOM performed the sequencing. GW did the aptamer structural prediction. All authors read and approved the final manuscript. COM, YMP, MJA, MSA, JLS, ASP, and YC wrote the manuscript.

## ACKNOWLEDGEMENT

The authors are grateful to the DNA sequencing platform at FIOCRUZ/RJ, and the National Center for Nuclear Magnetic Resonance (CNRMN) facility at UFRJ. This research used resources from the Brazilian Synchrotron Light Laboratory (LNLS), an open national facility operated by the Brazilian Center for Research in Energy and Materials (CNPEM) of the Brazilian Ministry for Science, Technology, Innovations and Communications (MCTIC). The SAXS1 beamline staff is acknowledged for the assistance during the experiments, in special Dr. Antônio Malfatti Gasperini. We thank MSc. Geisi R. Barreto for the assistance with the NTA measurements.

