## Supplementary data for "Liquid-liquid phase separation and aggregation of the prion protein globular domain modulated by a high-affinity DNA aptamer"

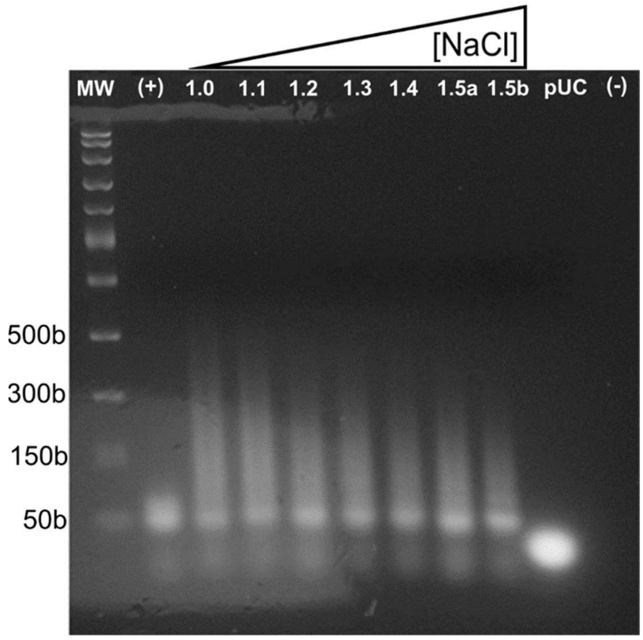


**Figure S1**. 2% Agarose gel electrophoresis of DNA aptamers selected against $rPrP^{90\text{-}231}$ eluted by NaCl gradient. 5 µL of eluted samples at different NaCl concentrations (from 1 to 1.5 M) identified as 1.0; 1.1; 1.2; 1.3; 1.4; 1.5a and 1.5b were analyzed. Aptamer library was used as positive control (+) and ultra-pure water as negative control (-) in the PCR reaction. MW ladder was Fast DNA ladder (Invitrogen, USA) and size (base numbers) is indicated at the left.

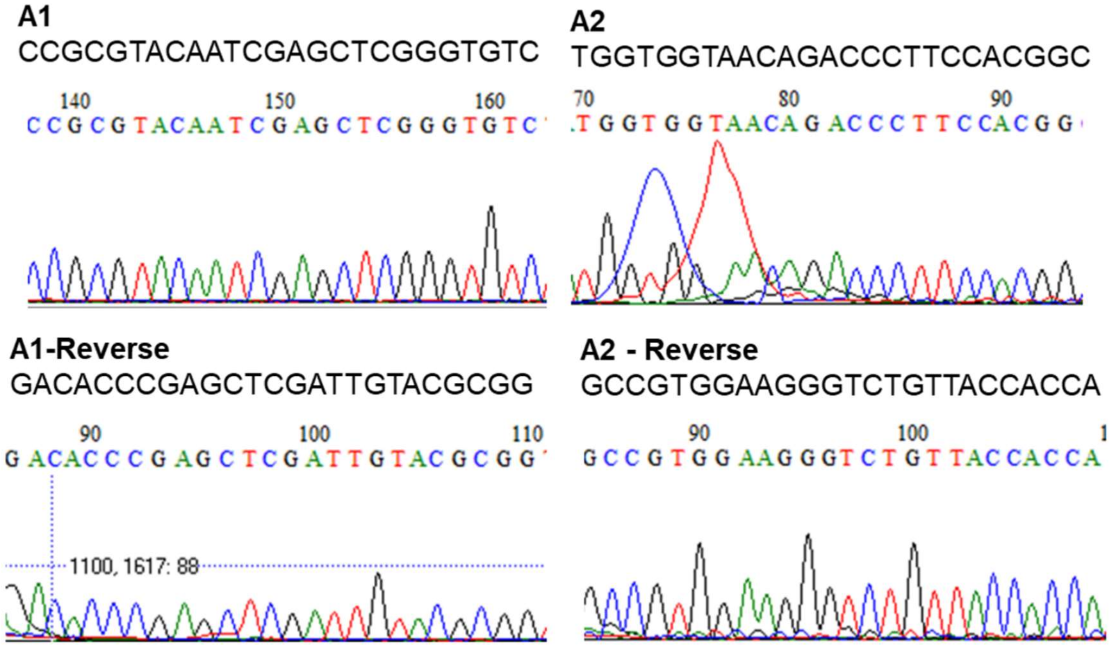


**Figure S2**. Electropherograms obtained from the sequencing analysis of isolated aptamers, using forward (top) and reverse (bottom) primers from the aptamer library. Only variable core region (25-mer) is shown, flanking regions were excluded.

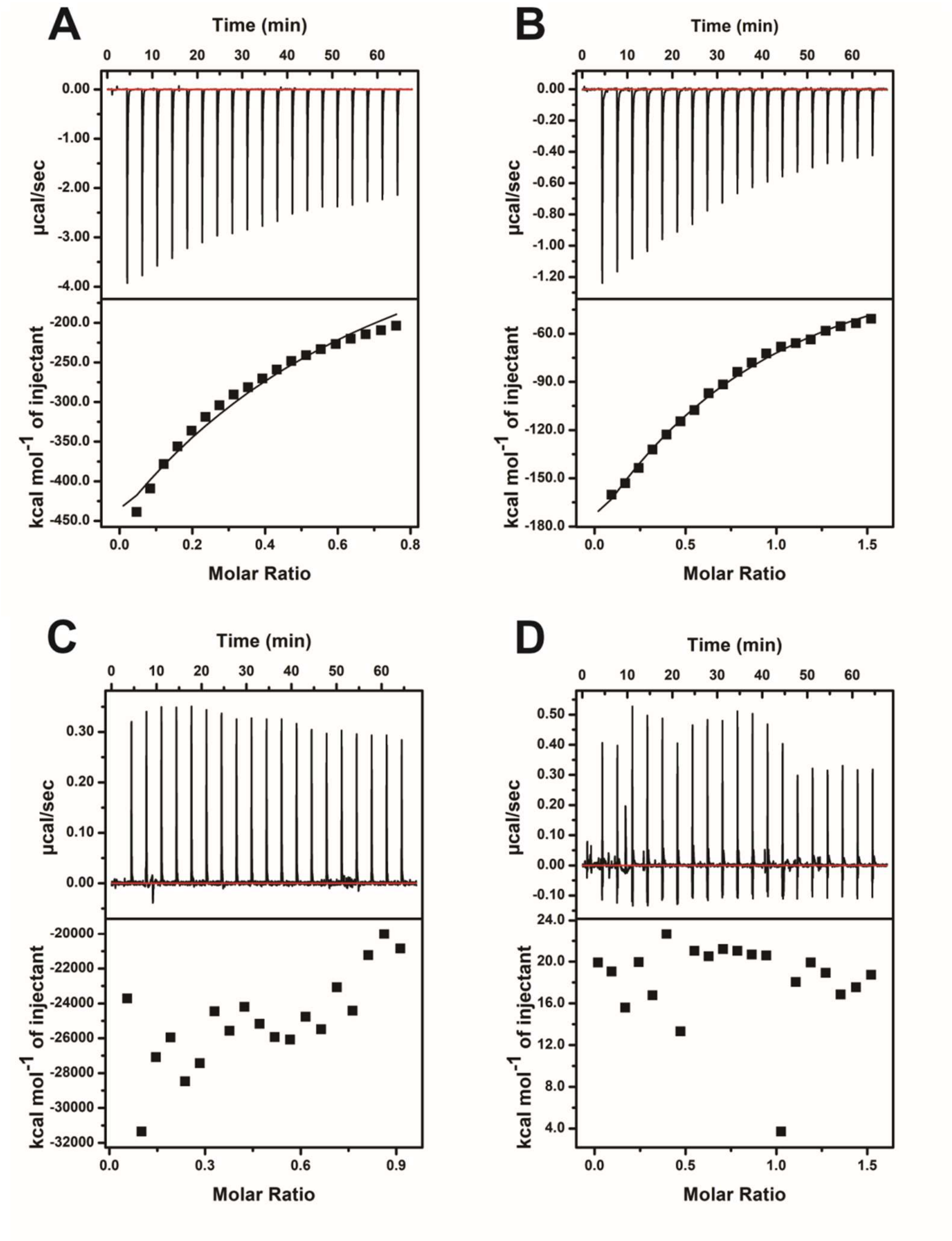


**Figure S3**. ITC binding isotherms of rPrP$^{90\text{-}231}$ with A1 and A2 in the presence of increasing salt concentrations. Raw ITC data (top panel) and the integrated heat values plotted against rPrP:aptamers (lower panel). (**A**) rPrP$^{90\text{-}231}$:A1 in 100 mM NaCl (**B**) rPrP$^{90\text{-}231}$:A1 in 500 mM NaCl (**C**) rPrP$^{90\text{-}231}$:A1 in 800 mM NaCl. (**D**) rPrP$^{90\text{-}231}$:A2 in 100 mM NaCl. Experiment was performed with 5 µM rPrP in 20 mM sodium cacodylate (pH 7.0) at 25°C.

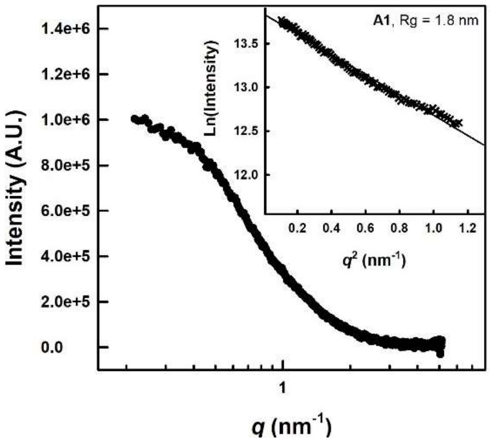


**Figure S4**. Small angle x-ray scattering curve of A1 and linear regression of the Guinier plot (inset) showing the obtained radius of gyration value.

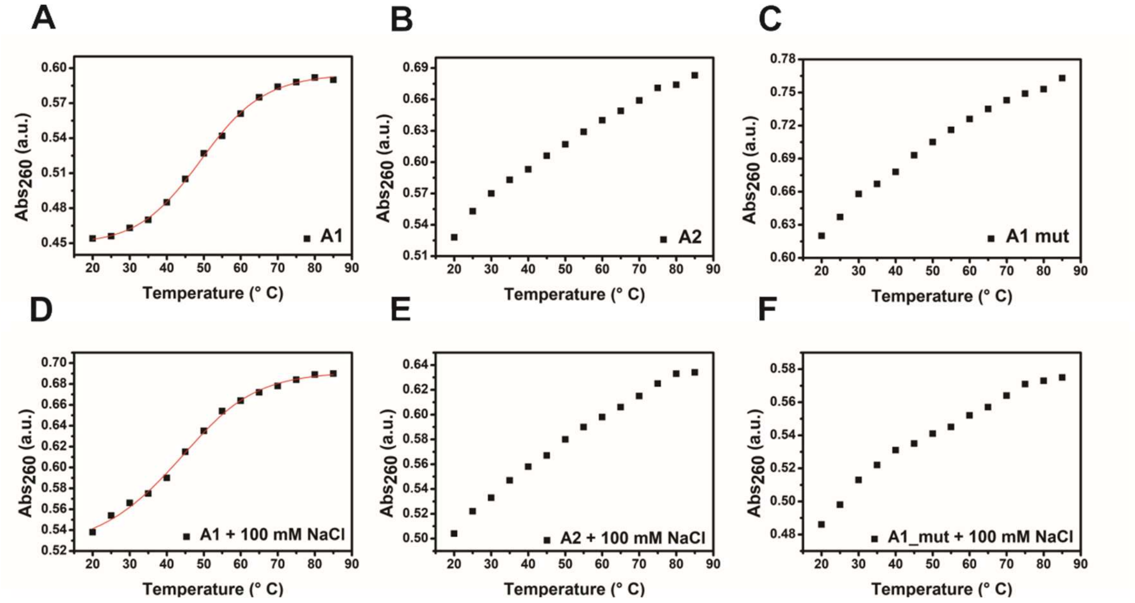


**Figure S5.** UV-Thermal denaturation profiles of A1 (**A**), A2 (**B**), and A1_mut (**C**) in the absence and presence (**D** to **F**) of 100 mM NaCl. Absorbance was measured at 260 nm and samples were heated from 20-85°C at a rate of 0.5°C /min. Sample concentrations were ~2 µM in 20 mM sodium cacodylate (pH 7.0).

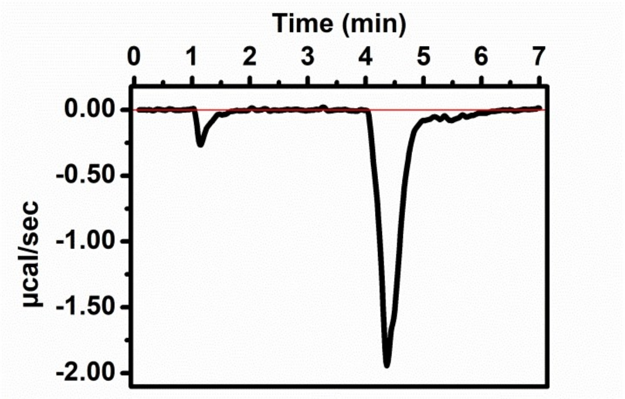


**Figure S6.** Single injection isothermal titration calorimetry of rPrP$^{90-231}$:A1_mut. Experiment was performed with 100 µM A1_mut in the syringe and 5 µM rPrP in 20 mM phosphate (pH 7.0), 25°C. Heat of dilution of A1_mut in buffer was measured separately and subtracted from the titration data. The final molar ratio was 1:1.

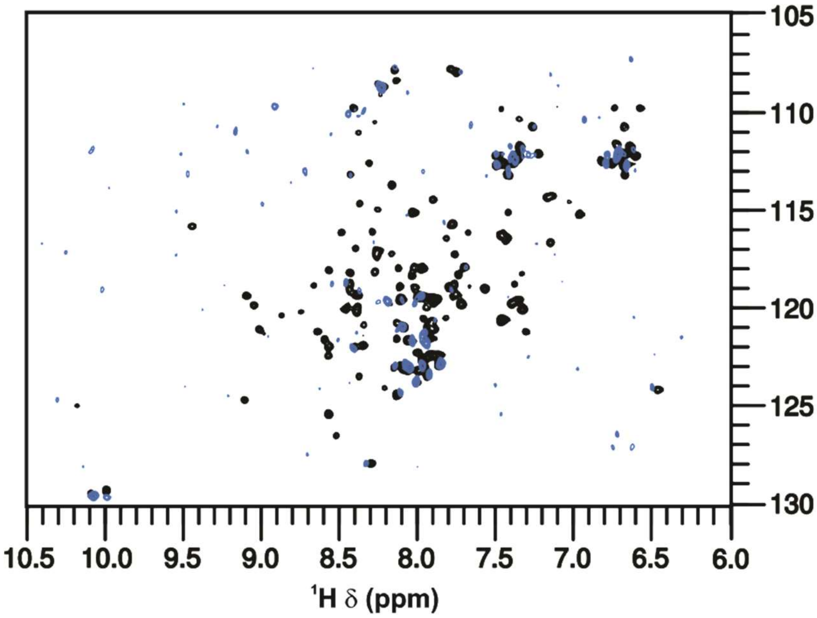


**Figure S7. NMR spectroscopy shows that rPrP$^{90-231}$ resonance signals disappear in the presence of A1.** Comparison of [$^{1}$H,$^{15}$N] HSQC spectra of 200 µM rPrP$^{90-231}$ acquired in the absence (black) and presence of 20 µM A1 (NA:protein, 1:10) (blue). NMR experiments were acquired in 10 mM potassium phosphate pH 6.5, 10 mM KCl at 25°C.

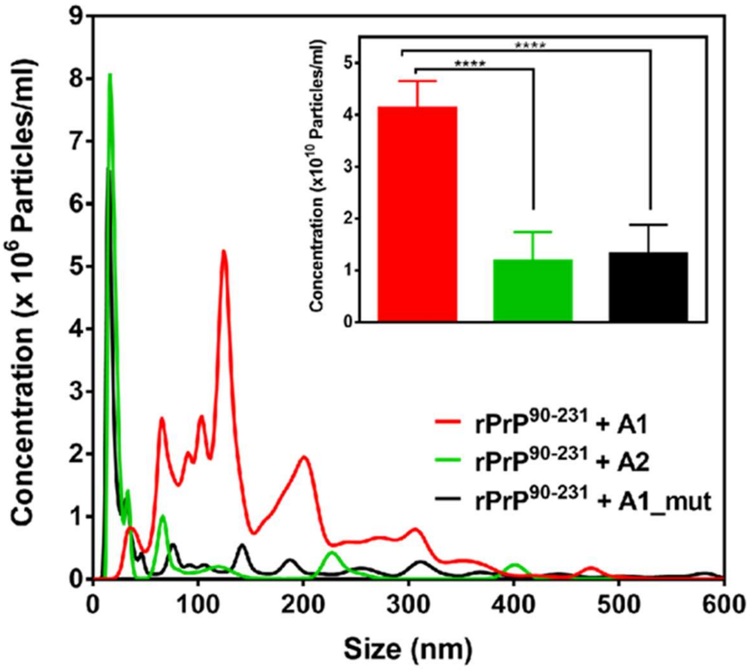


**Figure S8.** Particle size distribution and concentration of PrP:aptamer complexes assessed by Nanoparticle Tracking Analysis (NTA). Particle size distribution profile for $rPrP^{90\text{-}231}$ complexed with each aptamer (A1, A2 or A1_mut) at a 5:1 molar ratio. Lines represent the average of at least 4 readings per sample. Inset: Total concentration of nanoparticles in each sample. Error bars represent standard deviations, ****$P < 0.0001$. Readings were performed with 5 μM $rPrP^{90\text{-}231}$ and 1 μM aptamers in 10 mM Tris (pH 7.4), 100 mM NaCl at 25°C.

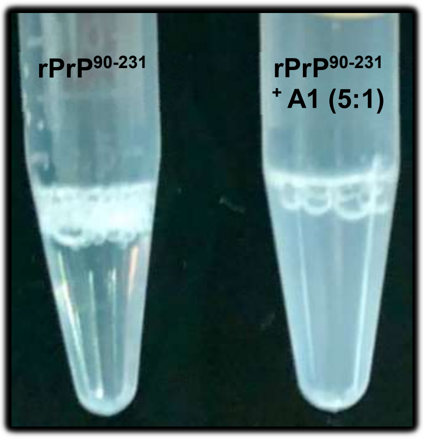


**Figure S9. $rPrP^{90\text{-}231}$ increased turbidity upon A1 aptamer incubation.** Upon addition of A1 aptamer at 5:1 (protein:aptamer) molar ratio instantly transition to a turbid solution occurs at room temperature, as visually observed on the right image.

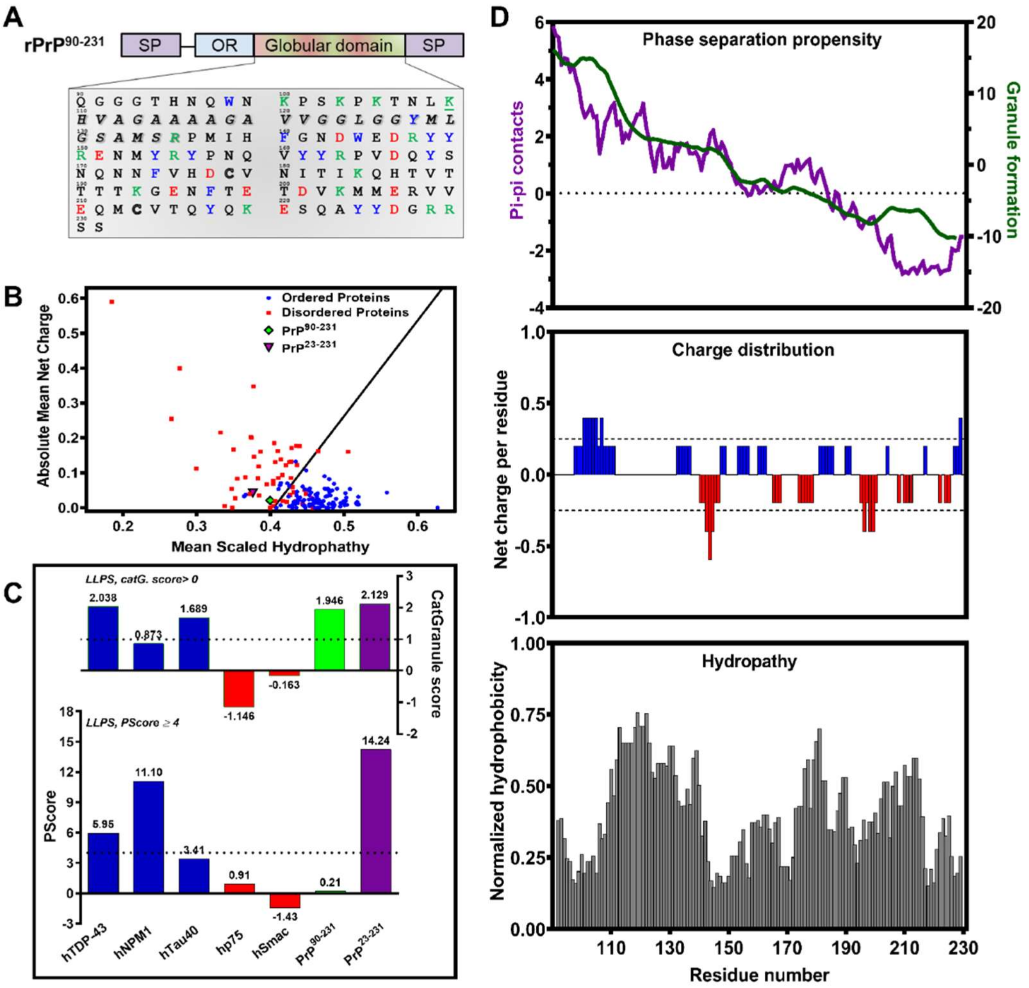


**Figure S10. Prediction of molecular determinants involved in rPrP$^{90\text{-}231}$ phase separation by bioinformatic tools**. (**A**) C-terminus of rPrP (residues 90 to 231) is enriched in positive (green), negative (red) and aromatic (blue) amino acids that can potentially interact through cation-pi, pi-pi, hydrophobic and electrostatic contacts. The hydrophobic region (highlighted in italic font) is a putative dimerization domain (45) and the two Cys residues, 178 and 213, (hollow font) are conserved. SP, signal peptide; OR, octapeptide repeat region. (**B**) PONDR® (Predictor of Natural Disordered Regions; http://www.pondr.com/) predictor analysis based on the charge-hydropathy plot revealed disordered nature of both full-length PrP and PrP$^{90\text{-}231}$. (**C**) Total propensity scores for LLPS based on catGranule (top) and pi-pi predictors (bottom). rPrP$^{90\text{-}231}$ (green bar) and rPrP$^{23\text{-}231}$ (purple bar) scores were compared with positive controls of proteins that undergo *in vitro*/*in vivo* phase transition (blue bars) or negative controls composed by folded proteins such as hSmac (Uniprot ID Q9NR28) and hp75 (Uniprot ID P08138) (red bars). (**D**) **Top:** pi-containing groups are found in known proteins that phase separate. The first segment of PrP$^{90\text{-}231}$, composed of approximately 84 residues, can interact through pi-pi contacts as screened by PScore (purple curve; significant total PScore ≥ 4; https://cdn.elifesciences.org/articles/31486/elife-31486-supp-v1.zip). CatGranule algorithm calculates granule-forming tendency based on primary sequence residues composition, structural disorder and nucleic acid binding property (green curve; score > 0 indicate proteins likely to undergo LLPS and scores > 1 are associated to strong ability to phase separate; https://tartaglialab.com/new_submission/catGRANULES). **Middle:** oppositely charged segments of $PrP^{90-231}$ evidenced in the plot of net charge along primary sequence using CIDER (Classification of Intrinsically Disordered Ensemble Regions, sliding window size of 5; http://pappulab.wustl.edu/CIDER). $PrP^{90-231}$ is a weak polyampholyte suggesting phase transition mediated by electrostatic interactions. **Bottom:** Hydropathy along sequence obtained from a normalized Kyte-Doolittle hydrophobicity ranging from 0 (lowest hydrophobicity) to 1 (highest hydrophobicity) using CIDER. Analyses shown in panel D show region from residues 90 to 231; the full-length protein sequence was used as input.

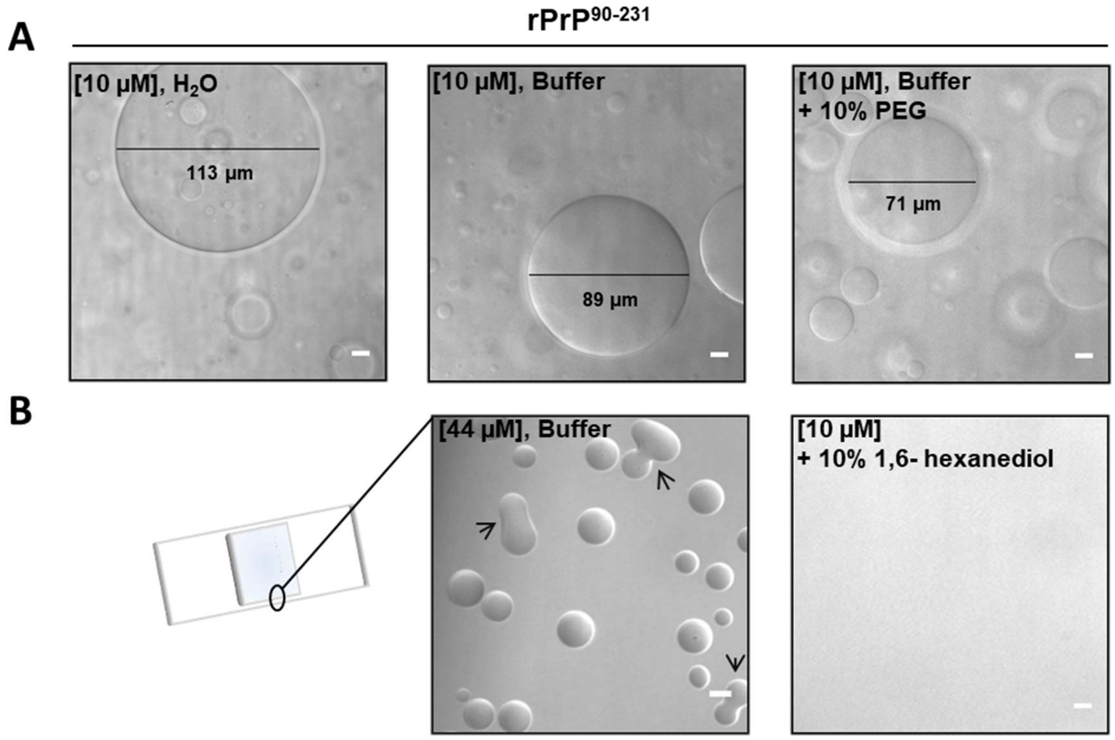


**Figure S11.** ***In vitro*** **rPrP$^{90\text{-}231}$ phase separation**. (**A**) rPrP$^{90\text{-}231}$ phase separates into droplets with varying sizes at low concentration (10 µM). Coalescence into large droplets occurs in water (left), buffer composed by 10 mM Tris pH 7.4, 100 mM NaCl (middle) and upon addition of 10% (w/v) PEG 4000 (right). The diameter is represented under black lines. (**B**) At the glass liquid-air boundary occurs protein supersaturation insofar as laser exposure causes evaporation. In this area, protein-rich rPrP$^{90\text{-}231}$ spherical condensates rapidly fuse with one another. The aliphatic alcohol 1,6-hexanediol at low concentration (10%) was able to disassemble liquid-like droplets since weak hydrophobic interactions are perturbed.

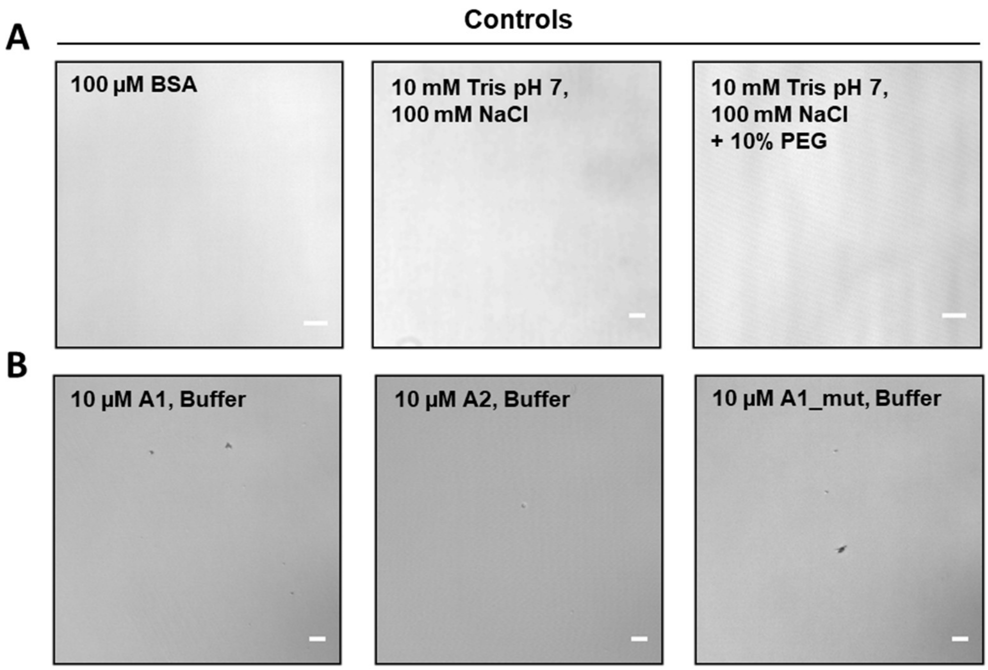


**Figure S12. DIC microscopy from controls of rPrP$^{90\text{-}231}$ phase separation assays.** (**A**) Bovine Serum Albumin (BSA) even at high concentration does not show liquid-like properties using the same vehicle for rPrP$^{90\text{-}231}$ assay (left). Buffer control (middle). Buffer in the presence of crowding agent, 10% PEG-4000 (right). (**B**) Micrographs from the highest concentration of aptamers (10 µM) used in the assays. A1 (left), A2 (middle) and A1_mut (right). Magnification, 630 ×. Scale bar, 10 µm.

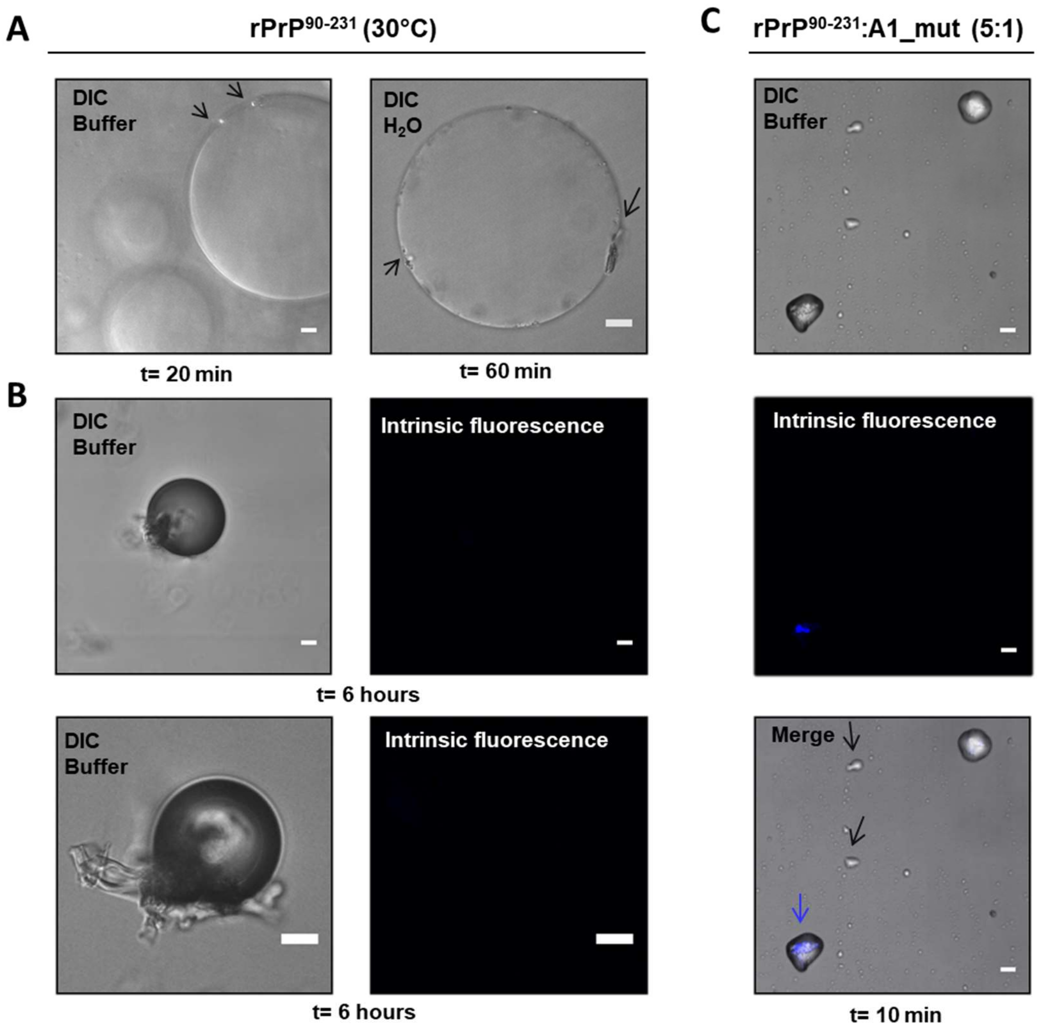


**Figure S13. rPrP$^{90-231}$ in the presence of A1_mut shows aberrant phase separation.** Representative images from DIC and intrinsic fluorescence suggestive of amyloid formation (excitation at 405 nm; emission at 450-500 nm). (**A**) Micrograph of 10 µM rPrP$^{90-231}$ in 10 mM Tris (pH 7), 100 mM NaCl (left) or in water (right) incubated at 30°C for specified times evidencing fiber-like material protruding from droplet surface (black arrows). (**B**) Same sample imaged after 6 hours does not show intrinsic fluorescence. Two representative images demonstrate dark retractile structures with uneven surface resembling aged droplets. Fiber-like material is more evident arising from the edges. These characteristics are compatible with an organized solid-like state. (**C**) rPrP$^{90-231}$ in the presence of A1_mut (5:1 molar ratio) in 10 mM Tris (pH 7), 100 mM NaCl. After 10 min incubation at 30°C, not perfectly spherical condensates were detected. Black arrows indicate "sticky droplets" that were not able to completely fuse. Intrinsic fluorescence arising from an "old" droplet surface is indicated by a blue arrow. Scale bar, 10 µm.

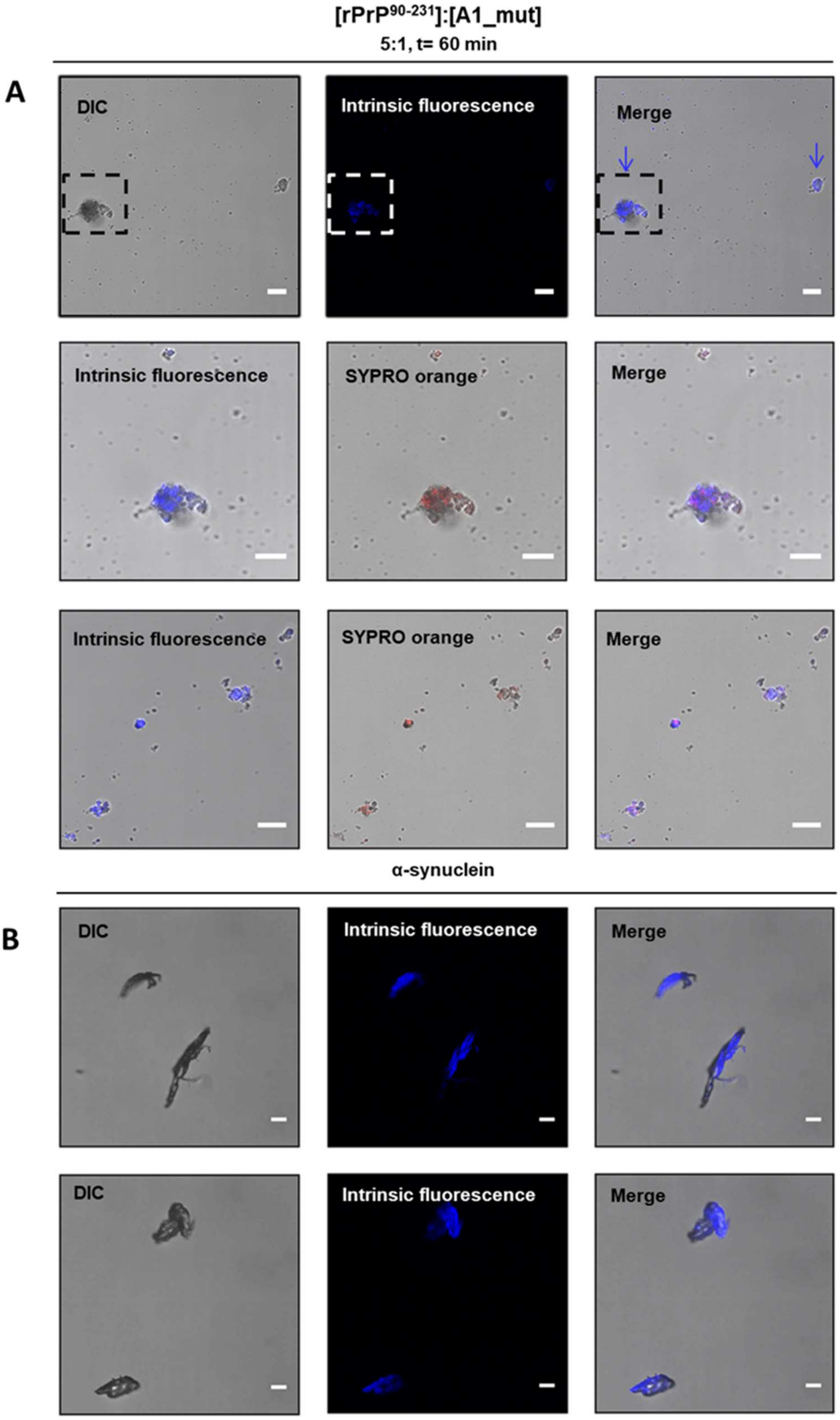


**Figure S14. rPrP$^{90-231}$ incubated with A1_mut forms aggregates that bind SYPRO orange and exhibit typical auto-fluorescence of amyloidogenic proteins.** DIC representative images, staining with SYPRO orange (488 nm excitation line; emission 500-650 nm) and intrinsic fluorescence suggestive of amyloids (excitation at 405 nm; emission at 450-500 nm). (**A**) Addition of A1_mut to rPrP$^{90-231}$ (1:5 molar ratio) in 10 mM Tris pH 7, 100 mM NaCl incubated for 60 min at room temperature**. Top:** Only solid-like aggregates with intrinsic fluorescence suggestive of amyloid structures were observed (blue arrows). **Middle:** Zoom of the top dashed square. Aggregate stained with SYPRO orange also exhibits intrinsic amyloid fluorescence. **Bottom:** Another microscopic region showing a significant number of SYPRO orange-positive aggregates possessing autofluorescence reminiscent of amyloidogenic proteins. (**B**) Autofluorescence positive control using α-synuclein fibrillar aggregates, prepared as described (43).
